# NRBP1 pseudokinase binds to and activates the WNK pathway in response to osmotic stress

**DOI:** 10.1101/2024.12.12.628181

**Authors:** Ramchandra V Amnekar, Toby Dite, Pawel Lis, Sebastian Bell, Fiona Brown, Clare Johnson, Stuart Wilkinson, Samantha Raggett, Mark Dorward, Mel Wightman, Thomas Macartney, Renata F Soares, Frederic Lamoliatte, Dario R Alessi

## Abstract

WNK family kinases are regulated by osmotic stress and control ion homeostasis by activating SPAK and OXSR1 kinases. Using a proximity ligation approach, we found that osmotic stress promotes the association of WNK1 with the NRBP1 pseudokinase and TSC22D2/4 adaptor proteins, results that are confirmed by immunoprecipitation and mass spectrometry and immunoblotting studies. NRBP1 pseudokinase is closely related to WNK isoforms and contains a RΦ-motif binding conserved C-terminal (CCT) domain, similar to the CCT domains in WNKs, SPAK and OXSR1. Knockdown or knock-out of NRBP1 markedly inhibited sorbitol-induced activation of WNK1 and downstream components. We demonstrate recombinant NRBP1 can directly induce the activation of WNK4 *in vitro*. AlphaFold-3 modelling predicts that WNK1, SPAK, NRBP1, and TSC22D4 form a complex, in which two TSC22D4 RΦ-motifs interact with the CCTL1 domain of WNK1 and the CCT domain of NRBP1. Our data indicates NRBP1 functions as an upstream activator of the WNK pathway.

**Teaser:** NRBP1 functions as a scaffolding component regulating the assembly of a multi-subunit complex, required for the activation of the WNK Lysine Deficient Protein Kinase family in response to osmotic stress.

*Graphical Abstract:* 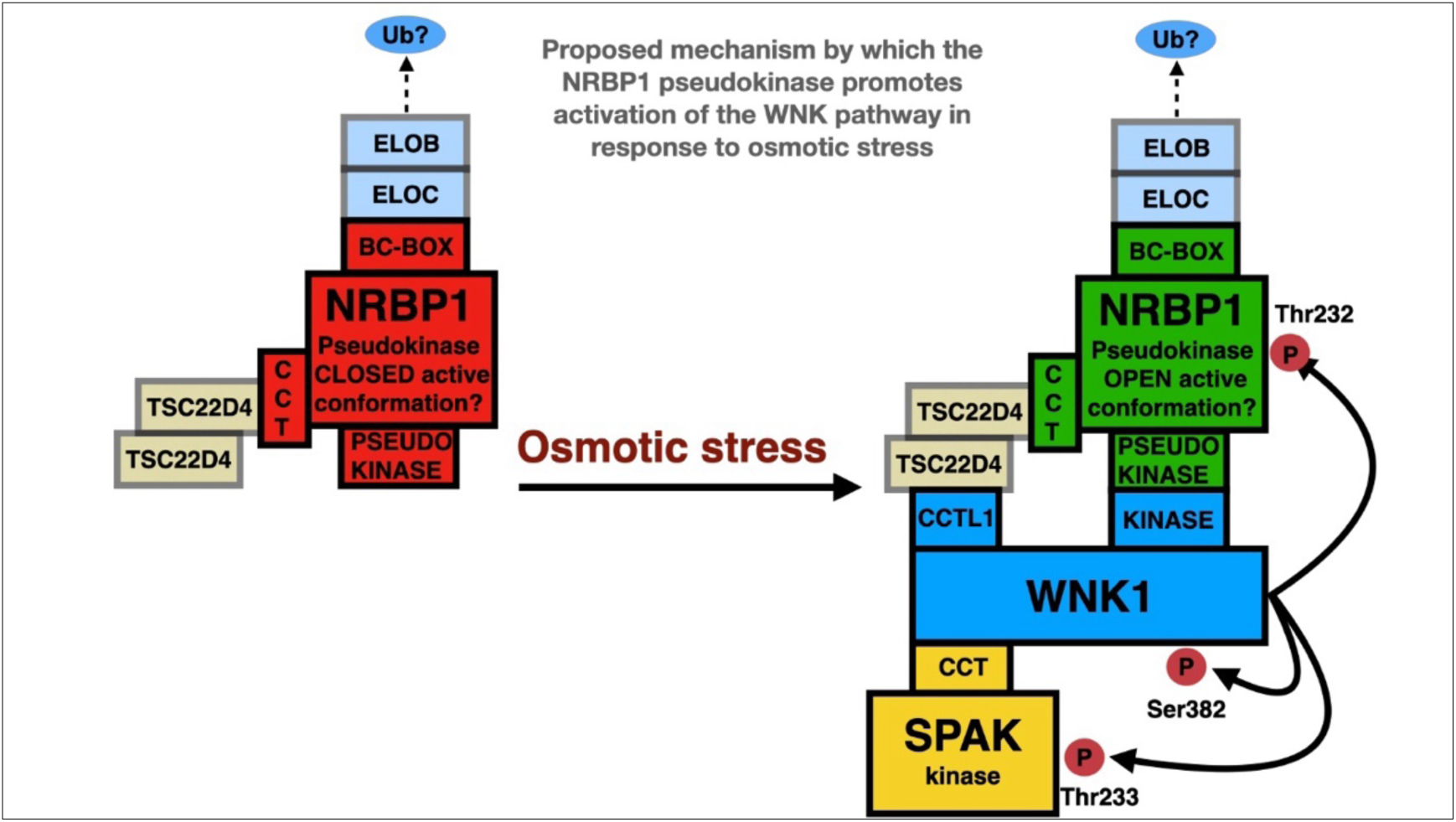

## Introduction

The With-No-Lysine (WNK) kinases are regulated by osmotic stress and chloride ions and are distinguished by their atypical catalytic lysine residue positioning (*1*). Under hypertonic stress or conditions of low intracellular chloride, WNK kinases undergo autoactivation through T-loop trans-autophosphorylation. This activation in turn leads to the phosphorylation of T-loop residues in two well-characterized downstream protein kinase substrates: STE20/SPS1-related proline-alanine-rich kinase (SPAK, also known as STK39) and oxidative stress responsive 1 kinase (OXSR1, previously known as OSR1) (*2–5*).

SPAK and OXSR1 contain a CCT (Conserved C-Terminal) docking domain that specifically interacts with conserved RΦXV-motifs (where Φ denotes a hydrophobic amino acid) on WNK isoforms, a process essential for their efficient activation (*4, 6–8*). Recent studies have also identified two CCT-like domains within WNK isoforms, designated CCTL1 (residues 481-571 on WNK1) and (residues 1115-1195 on WNK1) (*9–12*). While *in vitro* interactions have been observed between the CCTL1 domain of WNK1 and RΦ-motif peptides (*10, 11*), the physiological roles of these CCT-like domains in regulating the WNK signaling pathway remain unclear.

Key substrates of SPAK and OXSR1 include a group of electroneutral cation-chloride cotransporters that regulate intracellular chloride influx, cell volume, osmosensing, and blood pressure (*2, 13–17*). These cotransporters include the ubiquitously expressed NKCC1, as well as the kidney-specific NCC (Na–Cl cotransporter) and NKCC2 (Na–K–2Cl cotransporter 2). All of these are phosphorylated and activated by SPAK and OXSR1 (*4, 17–20*).NKCC1, NCC, and NKCC2 each contain conserved RΦ-motifs that bind to the CCT domain of SPAK and OXSR1. This interaction is essential for the efficient phosphorylation and activation of these ion cotransporters (*4, 6*). Thus, the CCT domain is crucial not only for the activation of SPAK/OXSR1 by WNK isoforms but also for the effective phosphorylation of their substrates.

Additionally, SPAK and OXSR1 also phosphorylate and inhibit the activity of four potassium-driven cation–chloride cotransporters (KCC1, KCC2, KCC3, and KCC4), which mediate chloride efflux from cells (*21–24*). The reciprocal regulation of electroneutral cation-chloride cotransporters by SPAK and OXSR1—activating chloride influx (NCC, NKCC1/2) and inhibiting chloride efflux (KCCs)—ensures precise control of intracellular chloride levels (*2, 25*).

At the organismal level, the WNK-SPAK/OXSR1-cation chloride cotransporter pathway is crucial for regulating ion reabsorption in the kidney, which plays a vital role in controlling blood pressure (*13, 17, 26, 27*). Mutations that lead to increased levels of WNK1 or WNK4 in the kidney are associated with rare forms of hypertension (*28, 29*). Recent research has also shown that WNK kinases contribute to a variety of other cellular processes, including autophagy (*30*), cancer (*31, 32*), neuronal development (*33*), mitosis (*34*), T cell adhesion and migration (*35*), as well as inflammasome activation and pyroptosis (*36, 37*).

Given the broad roles of WNK isoforms in regulating a variety of cellular processes, we hypothesized that additional key regulators of the WNK signaling pathway likely exist. Early co-immunoprecipitation studies of WNK1 led to the discovery of its key substrates, SPAK and OXSR1 (*3*), and provided evidence that WNK isoforms can form homo- and heterodimers (*38*). Co-immunoprecipitation experiments also confirmed genetic findings that WNK4 interacts with the KLHL3 and CUL3 E3 ligase components, which regulate intracellular WNK4 levels (*39*). Although several other proteins, including LINGO-1, SGK1, TAK1, and EMC2, have been reported as WNK1 interactors (*40–43*), the functional significance of these remains unclear.

In this study, we employed a proximity labeling approach to identify novel regulators of the WNK1 pathway, a method that has proven successful in uncovering arrays of functional regulators in other signaling pathways (*44, 45*). Using this strategy, we identified a pseudokinase Nuclear Receptor Binding Protein 1 or NRBP1, which is closely related to WNK isoforms, as a novel interactor. Our studies reveal that NRBP1 functions as a critical regulator of WNK1 signaling pathway.

## RESULTS

### Osmotic stress induces interaction between WNK1 and NRBP1 pseudokinase

To identify novel WNK interactors we employed a proximity ligation approach (*44, 45*). We generated N-terminal GFP tagged WNK1 knock-in human embryonic kidney (HEK293) cell lines stably expressing the Flag Turbo-ID biotin ligase fused to αGFP6M, an anti-GFP nanobody. This approach is designed to bring the Turbo-ID biotin ligase to GFP-WNK1 and biotinylate proximal proteins (Fig 1A) and has been utilized previously to identify potential substrates of the CUL2 E3 ligase diGly receptor KLHDC2 (*46*). We observed that treating GFP-WNK1 knock-in HEK293 cells expressing Flag TurboID-αGFP6M with exogenous 0.5 mM biotin for 5 min, prior to cell lysis, markedly boosted biotinylation of proteins including one corresponding to the molecular weight of GFP-WNK1 (Fig 1B). These data are consistent with previous work showing that exogenous biotin improves efficiency of TurboID mediated biotinylation (*45*). We next sought to exploit this system to identify proteins that are in proximity to WNK1 in an osmotic-stress-dependent manner. We treated wild type and GFP-WNK1 knock-in cells expressing Flag TurboID-αGFP6M with 0.5 mM biotin in the presence or absence of 0.5 M sorbitol for 5 min and biotinylated proteins were enriched by streptavidin affinity purification and analyzed by immunoblot analysis (SFig 1A) and DIA mass spectrometry (Fig 1C, SFig 1B).

**Figure 1:**
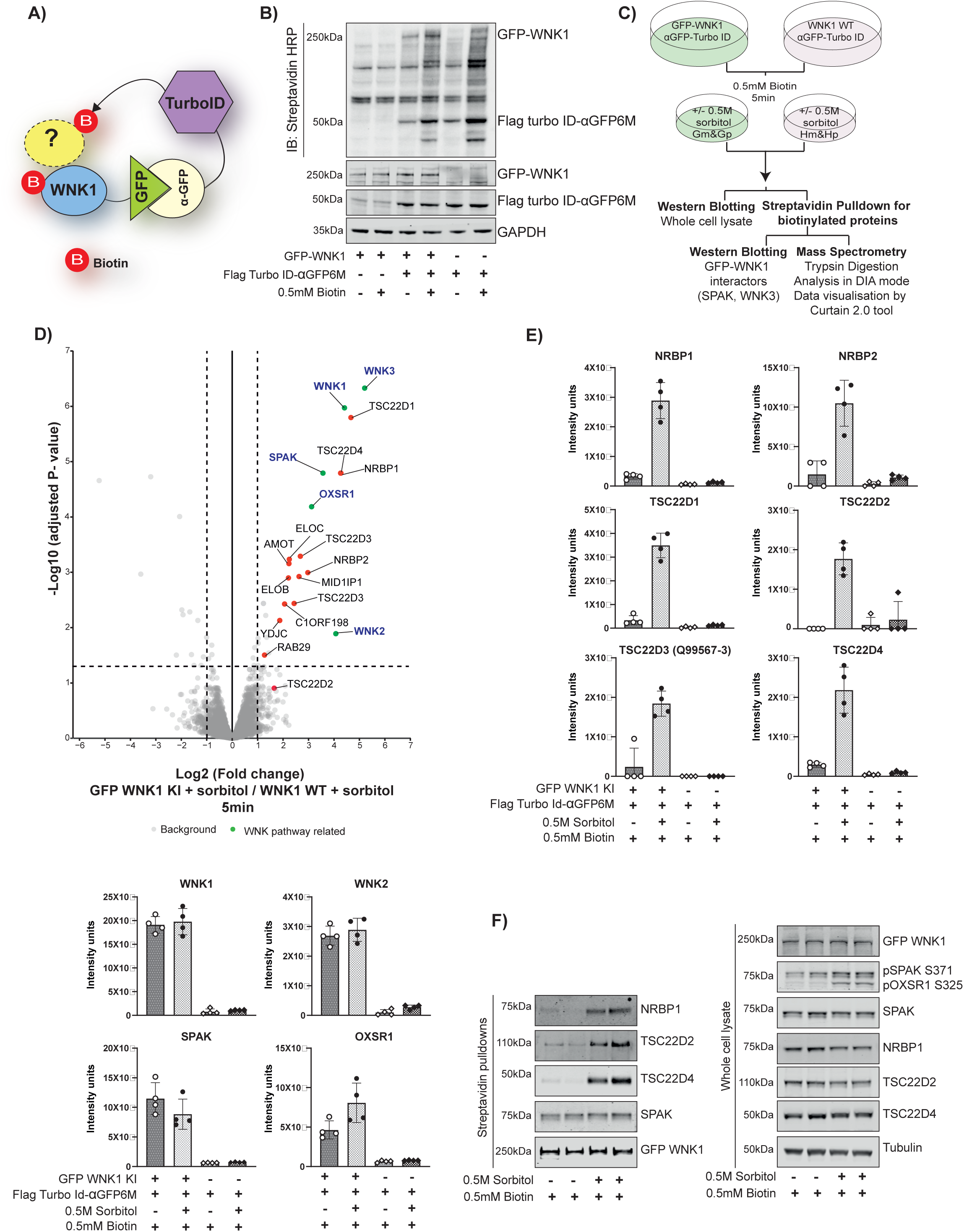
WNK1 associates with NRBP1, TSC22D2 and TSC22D4 during hypertonic stress. **(A):** Schematic of the GFP-WNK1 + FLAG-TurboID-aGFP6M system. TurboID utilizes biotin and ATP to produce biotinyl-AMP, which covalently biotinylates proximal nucleophilic amino acids (e.g., lysine) on proteins adjacent with WNK1. **(B)** Immunoblots of the indicated cell extracts with or without FLAG-TurboID-aGFP6M expression and/or exogenous biotin treatment. **(C)** Schematic of the TurboID-MS experiment under hypertonic stress. GFP-WNK1 knock-in (KI) and wild-type (WT) HEK293 cells expressing FLAG-TurboID-aGFP6M were treated with or without 0.5 M sorbitol for 5 minutes in the presence of 0.5 mM biotin (n=5 per treatment group). Streptavidin pulldown was followed by washing, S-trap microcolumn peptide preparation, and MS analysis in DIA mode. Data was processed in Python and visualized using the Curtain tool. Note that G-denotes GFP WNK1 KI cells and H-denotes WT WNK1 cells; m-no sorbitol and p-with sorbitol. **(D) Top Panel:** Volcano plots showing proteins enriched in GFP-WNK1 KI HEK293 cells compared to WT HEK293 cells under sorbitol treatment. Proteins highlighted in green include WNK1 and its known interactors (e.g., WNK2, WNK3, SPAK, OXSR1). Proteins in red represent sorbitol-specific interactors of WNK1. The plots show proteins with ≥2-fold enrichment and statistical significance, normalized to the median intensity of all proteins, with missing values imputed using a Gaussian distribution. P-values were adjusted using the Benjamini-Hochberg method, with a significance threshold of corrected P < 0.05. Data are based on five technical replicates per group. **Bottom panel**: Box plots showing the median intensities of significant hits identified in the volcano plot. Curtain-link-https://curtain.proteo.info//84e40fb8-144e-4859-9b2b-924091194344. **(E)** Box plots of median protein intensities for WNK1 interactors enriched specifically under hypertonic stress conditions. **(F)** Lysates and streptavidin pull-downs from GFP-WNK1 TurboID cells confirm biotinylation of NRBP1, TSC22D2, and TSC22D4, specifically following hypertonic stress.

The phosphorylation of SPAK (pSPAK-S371) and OXSR1 (pOXSR1-S325) confirmed the activation of WNK pathway. We found that WNK pathway components WNK1, WNK2, WNK3, SPAK and OXSR1 are biotinylated similarly in the absence or presence of sorbitol (Fig 1D, SFig 1C-E). We focused our attention on three interactors that are markedly induced by sorbitol, namely the pseudokinase NRBP1 (enriched 9.7-fold), and two of its known interactors namely transforming growth factor β-stimulated clone 22 domain family 2 and 4 (TSC22D2 enriched 3.2-fold and TSC22D4 enriched 6.869 fold) (*47–49*).Other TSC22D family members namely TSC22D1 (9.8-fold) and TSC22D3 (8-fold) isoforms were also found to be enriched along with ELOB (5.97 fold) and ELOC (6.72 fold) which are also known NRBP1 interactors (*48*) (Fig 1D-E, SFig1C-E). A recent study has also reported co-association of NRBP1, TSC22D family proteins and WNK1 in biomolecular condensates (*50*). Another recent study that we are co-authors also reports that NRBP1 interacts with the WNK4 isoform of WNK family (*51*). Previous work has shown that NRBP1 comprises a pseudokinase and is devoid of the ability to catalyze phosphorylation as it lacks three critical motifs in its pseudokinase domain that are essential for kinase activity (DFG, HRD and VAIK) (*52, 53*). The increased interaction of NRBP1, TSC22D2 and TSC22D4 with GFP1-WNK1 following sorbitol stimulation was also validated by independent immunoprecipitation and immunoblotting studies (Fig 1F).

### Domain structures of NRBP1, TSC22D2 and TSC22D4

The pseudokinase domain of NRBP1 is phylogenetically closely related to WNK1 (*54, 55*), and potentially NRBP1 could have been termed WNK5 (Fig 2A). Earlier work has also indicated that TSC22D2 and TSC22D4 play a role in protection from hypertonic stress (*56*). Furthermore, the DepMap database suggests that there is a significant codependence in cell viability between WNK1, NRBP1 and TSC22D2 in certain cancer cell lines, consistent with these components operating within a common pathway (*57, 58*) and this was also highlighted in another recent study (*50*).

**Figure 2:**
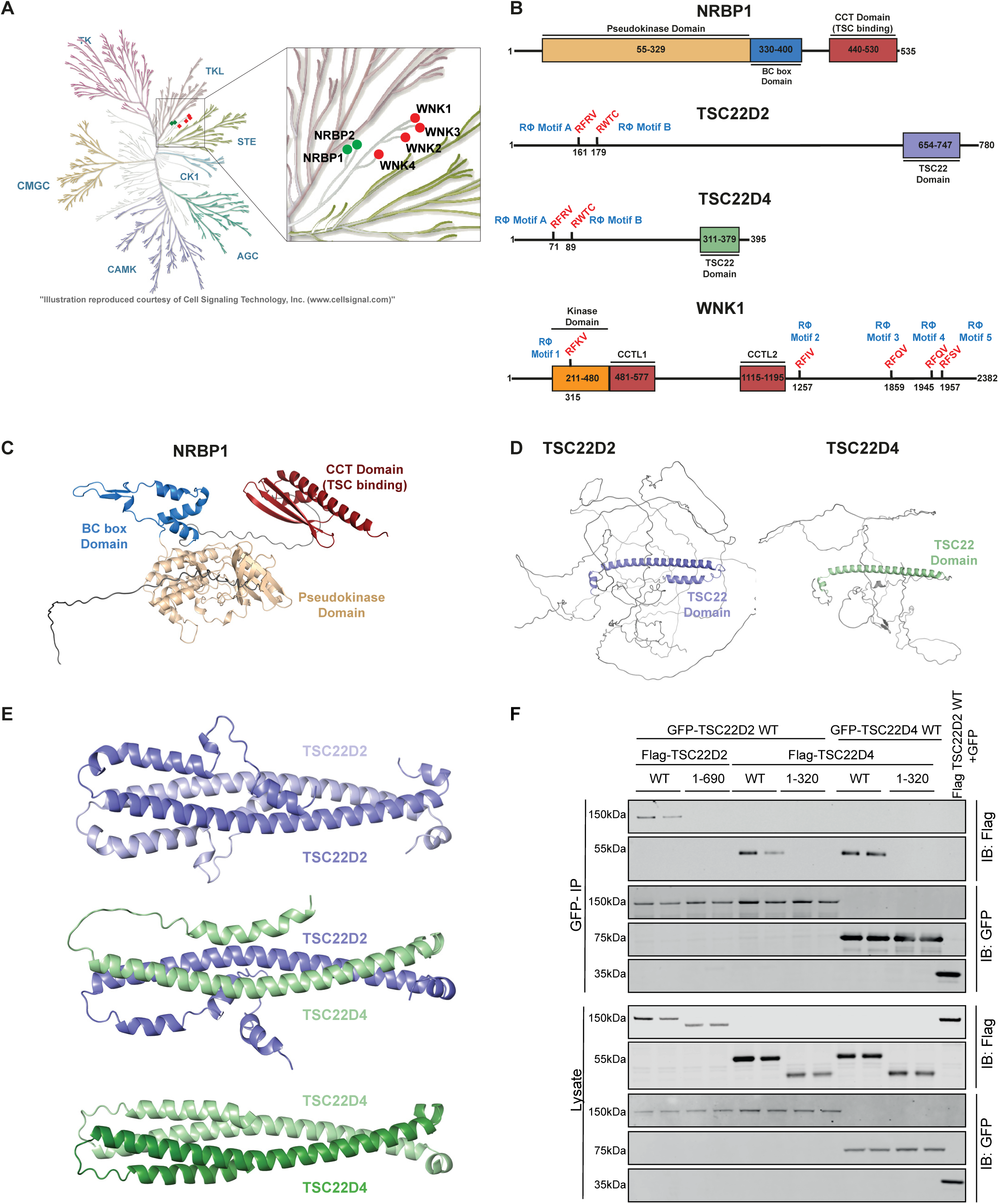
Domain organization of NRBP1, TSC22D2 and TSC22D4. **(A)** Phylogenetic Tree displaying evolutionary relationship between WNK kinases (highlighted in red) and the pseudokinases NRBP1 and NRBP2 (highlighted in green). **(B)** Schematic representation of the domain organization of human NRBP1, TSC22D2, TSC22D4, and WNK1 proteins. Key domains include the TSC-TGFβ1-stimulated clone 22 domain (TSC-TGFβ1), BC box (Elongin B and C binding domain), Src homology (SH) domain, and conserved C-terminal domains (CCT). The Rϕ motifs (critical for interaction) are highlighted in red, with two motifs in TSC22D2/4 and five in WNK1. **(C&D)** Predicted structural models of full-length NRBP1 and TSC22D2/4 proteins generated using AlphaFold3. **(E)** AlphaFold3-predicted structures of TSC22D2/4 homo- and heterodimers, focusing on the highly structured C-terminal TSC22 domain. Only the TSC22 domains are depicted, showing multiple contact sites critical for dimerization. **(F)** Co-immunoprecipitation assay in HEK293 cells co-transfected with GFP-TSC22D2/4 full-length (FL) wild-type and C-terminal deletion mutants of FLAG-TSC22D2. TSC22D2Δ refers to the TSC22D2 mutant lacking residues C-terminal to N690, while TSC22D4Δ lacks residues C-terminal to N320. Immunoprecipitation and immunoblotting confirmed homo- and heterodimerization of the full-length and mutant proteins. N=2 with two technical replicates each.

NRBP1 (535 residues) possesses a pseudokinase domain at its N-terminus, followed by a BC-box binding domain known to interact with elongin B and elongin C, components of the Cullin 2 and Cullin 4A E3 ligase ubiquitin system, and a C-terminal globular domain (*49, 59*) (Fig 2B, 2C). Alphafold-3 (AF3) (*60*) tertiary structure prediction analysis indicates that the C-terminal domain of NRBP1 is similar to the conserved C-terminal domains (CCT) found on WNK as well as SPAK and OXSR1 isoforms (Fig 2C, SFig 2A). This observation was also noted in another recent study (*61*). The AF3 predicted tertiary structure of TSC22D2 (780 residues) and TSC22D4 (395 residues) suggests that most of protein is disordered, apart from a long α helix region lying towards the C-terminus, encompassing a region of these proteins referred to as the TSC22 domain. The N-terminal of TSC22D2/4 proteins harbors two highly conserved potential RΦXX CCT binding motifs (*47*), designated as RΦ-motif in our study (Fig 2B, 2D). We have termed these as RΦ-motif-A (RFRV) and RΦ-motif-B (RWTC). It should be noted that RΦ-motif-B (RWTC) possesses a Trp residue rather than a Phe residue present in the canonical RFXV motif. Previous work has suggested that the C-terminal domain regions of TSC22D2 and TSC22D4 form a coiled-coil homo and hetero dimerization domain (*62, 63*), which is supported by AF3 modeling (*64*) (Fig 2E). Consistent with this, we observed that deletion of the TSC22 C-terminal domain of either TSC22D2 or TSC22D4 ablated co-immunoprecipitation with full length proteins (Fig 2F).

### Characterizing interaction of NRBP1-CCT domain with TSC22D2/4

We next explored how NRBP1 might interact with TSC22D2 and TSC22D4 using Alpha Fold-3 analysis (*60*). The data suggested that highly conserved residues of TSC22D2 (158 to 184) (Fig 3A) and TSC22D4 (70 to 94) (SFig 3A) encompassing the RΦ-motif-A (RFRV) and RΦ-motif-B (RWTC) discussed above, could interact with the NRBP1-CCT domain (Fig 3B). AF3 modeling indicates that the non-canonical RΦ-motif-B (RWTC) in both TSC22D2 (Fig 3A) and TSC22D4 (SFig 3A) bind to the CCT domain of NRBP1. To further explore this, we mutated the RΦ-motif-A and RΦ-motif-B motifs of TSC22D2 (Fig 3C) and TSC22D4 and tested impact on NRBP1 binding in a HEK293 cell co-expression system. For TSC22D2, consistent with the AF3 modeling, mutation of the RΦ-motif-B (R179A and R179E mutation) markedly impacted binding to NRBP1 whereas in case of RΦ-motif-A mutants the binding was reduced to half with the R161A+F162A mutant. (Fig 3C). Previous work undertaken in *Drosophila* demonstrated that fragments of TSC22D2 and TSC22D4 encompassing both RΦ-motifs motifs, co-immunoprecipitated with a C-terminal region of NRBP1 encompassing the CCT domain (*47*). To further validate the mechanism by which NRBP1 binds to TSC22D2 and TSC22D4, we also mutated two NRBP1 CCT domain residues (E484A, E492A) predicted by AF3 to interact with the Arg residue of the RΦ-motif-2 of TSC22D2 and TSC22D4 (Fig 3B, SFig 3A). The ability of the NRBP1[E484A+E492A] mutant to co-immunoprecipitate TSC22D2 and TSC22D4 was markedly reduced compared to wild type NRBP1 (Fig 3D, SFig 3C).

**Figure 3:**
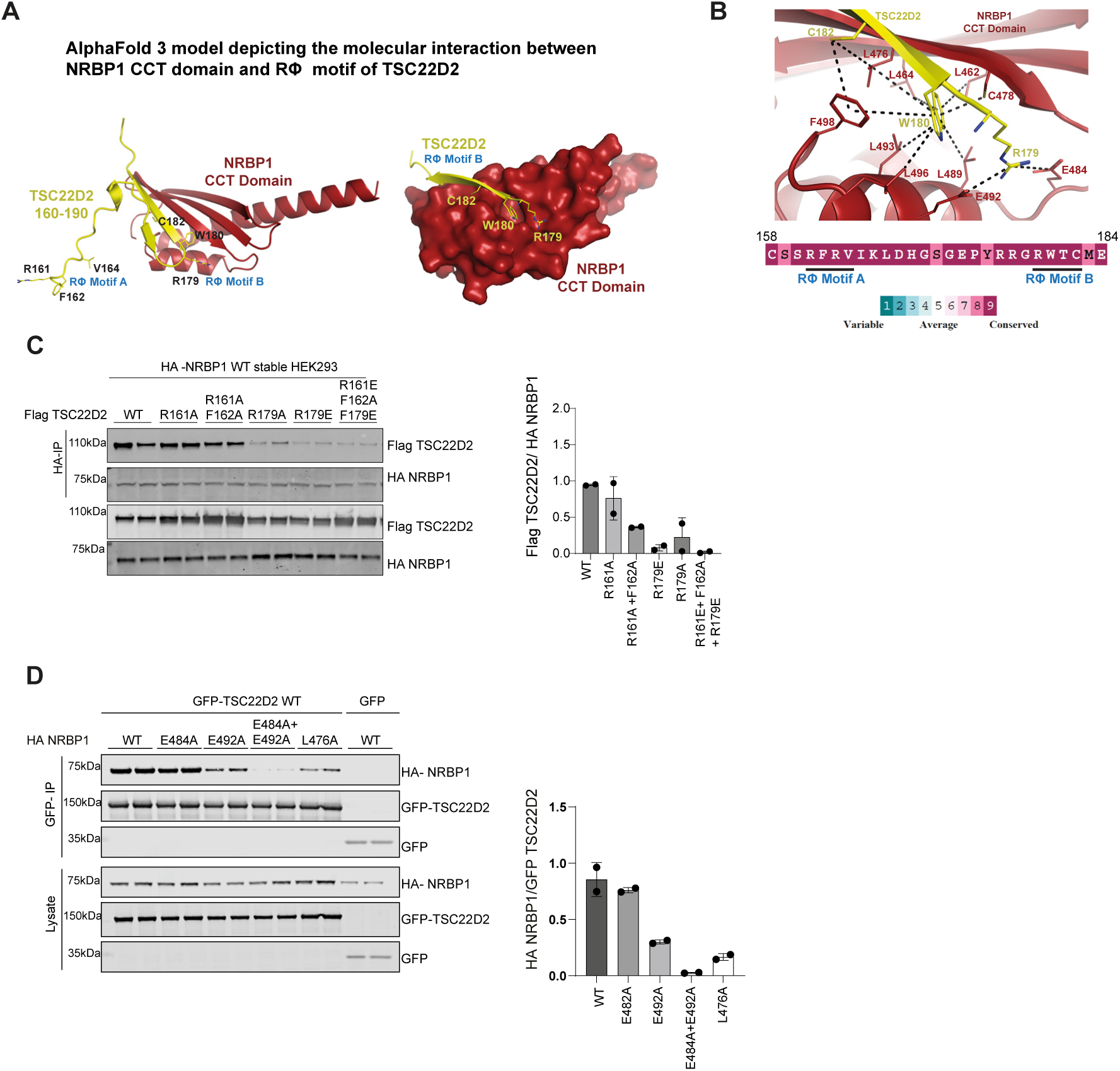
CCT like domain of NRBP1 interacts with the RϕXX motifs of TSC22D2 and TSC22D4. Structural modeling using AlphaFold3 reveals the molecular interaction between TSC22D2 and NRBP1. Residues 158–184 of TSC22D2, containing RΦ motif B, directly interact with NRBP1’s CCT-like domain. **(B)** Two glutamate residues (Glu484 and Glu492) in the CCT-like domain of NRBP1 form salt bridges with Arg179 of TSC22D2’s motif B. Additionally, Trp180 of TSC22D2 embeds into a hydrophobic pocket of NRBP1’s CCT domain, involving residues Leu462, Leu464, Leu476, Cys478, Leu489, Leu493, Leu496, and Phe498. Cys182 of TSC22D2 further stabilizes this interaction via hydrophobic interactions with Leu476 and Phe498 of NRBP1. Conservation analysis using ConSurf (*80*), demonstrates high evolutionary conservation of RΦ motifs A and B, underscoring their functional significance. **(C)** Left-HEK293 cells stably expressing HA-NRBP1 were transfected with FLAG-TSC22D2 mutants harboring alterations in RΦ motifs A and B. After 36 h, HA immunoprecipitation was performed to assess the interaction between NRBP1 and TSC22D2 mutants. FLAG-TSC22D2 levels in the HA-IP fraction were normalized to those in whole-cell lysates for analysis. Right-Densitometric analysis of the immunoprecipitated samples from two independent experiments, each performed in duplicate.

### WNK1 phosphorylates NRBP1 on its T-loop residue

We next explored whether recombinant WNK1 could phosphorylate NRBP1. We incubated recombinant wild type and kinase inactive (WNK1 V403F mutant) WNK1 (residues 1-661) with either full length NRBP1 or kinase inactive OXSR1(D164A) (all proteins expressed in *E. coli*) in an *in vitro* Mg[γ-^32^P]ATP phosphorylation reaction for 30 and 60 min and monitored phosphorylation following ^32^P-autoradiography of a Coomassie stained gel. We observed that wild type WNK1 phosphorylated NRBP1 to a similar extent as kinase inactive OXSR1, a physiological substrate of WNK1 (Fig 4A). No phosphorylation of NRBP1 or OXSR1 was observed with kinase inactive WNK1. Our data also suggested that the autophosphorylation of WNK1 was significantly enhanced by NRBP1 (Fig 4A), and this is discussed further in Figure 7.Trypsin digestion and mass spectrometry analysis of the GST-NRBP1 revealed a phosphorylated peptide encompassing the T-loop of the pseudokinase domain, phosphorylated at Thr-232 which was not observed with the kinase dead WNK1 (Fig 4B). A ClustalW sequence alignment indicates that the T-loop NRBP1-Thr232 residue is not conserved in WNK1, SPAK or OXSR1 and lies three residues before the WNK1 (Ser382), SPAK (Thr231) or OXSR1 (Thr185) T-loop sites (Fig 4C). The Thr232 site in NRBP1 is conserved through evolution from mammals to frog and fish, but not in Fly (SFig 4A). We generated a sheep polyclonal phospho NRBP1-Thr232 antibody which specifically detected wild type NRBP1 but not NRBP1[Thr232A] in an *in vitro* kinase assay with WNK1 and Mg-ATP (Fig 4D).

**Figure 4:**
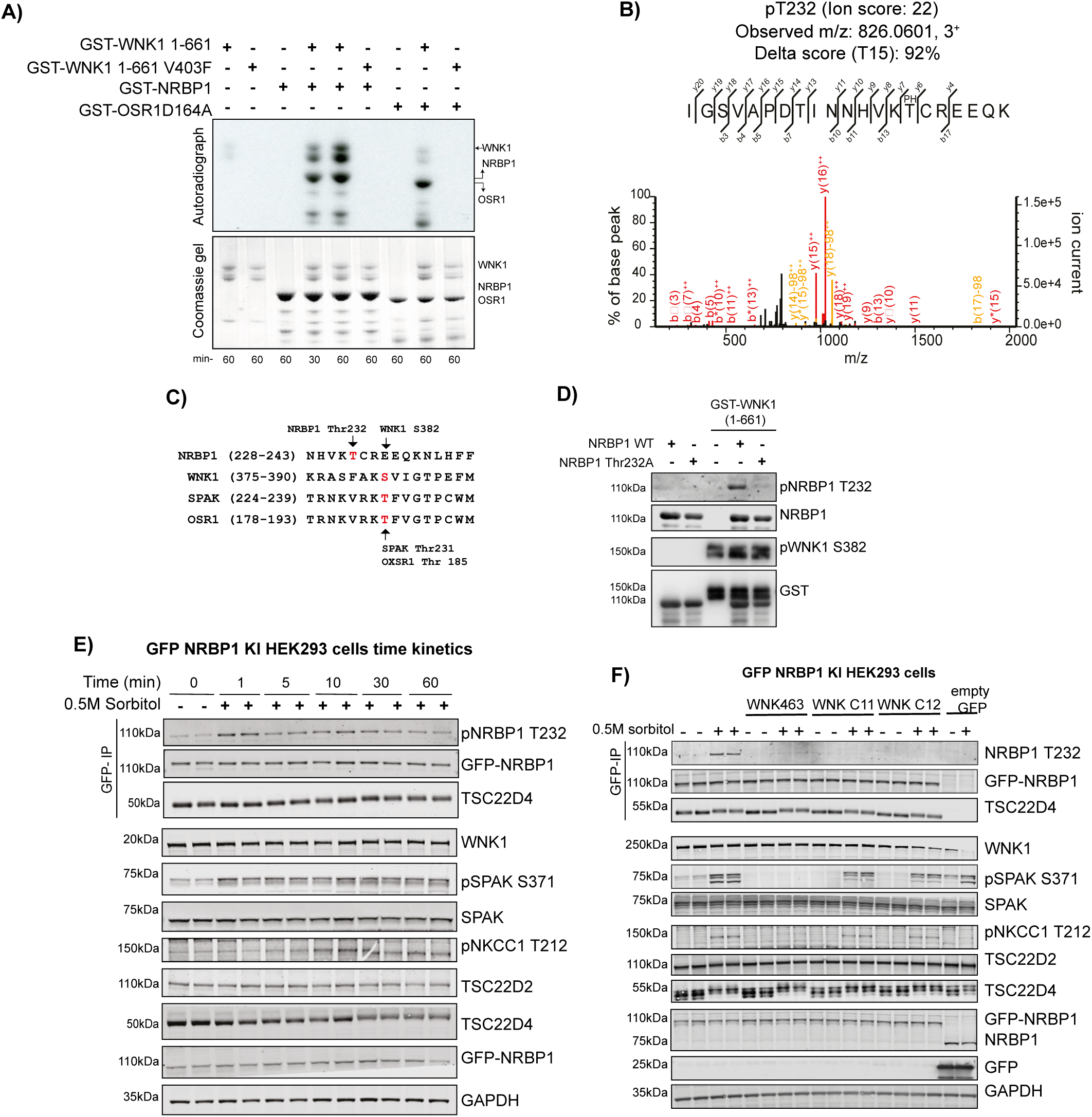
WNK1 directly phosphorylates NRBP1 *in vitro* and in cells. **(A)** Autoradiograph (top) and Coomassie-stained gel (bottom) showing phosphorylation of NRBP1 by the WNK1 kinase domain (residues 1–661) in an *in vitro* kinase assay. **(B)** MS/MS spectra confirming phosphorylation of the NRBP1 peptide at Thr232, the putative T-loop phospho-acceptor residue. Data are from three technical replicates for each condition. **(C)** Protein sequence alignment of T-loops from NRBP1, SPAK (STK39), and OXSR1 (OXR1), highlighting Thr232 of NRBP1 as a conserved phosphosite. The alignment was performed using MUSCLE (*81*) and visualized with JalView (*82*), demonstrating motif similarity to other known T-loop phosphorylation sites. **(D)** Immunoblot analysis using a sheep phosphospecific NRBP1 Thr232 antibody confirms phosphorylation in a kinase assay with GST-tagged NRBP1 (WT) and a Thr232A mutant by recombinant GST-WNK1 kinase domain (2–661). **(E)** Immunoblot analysis of GFP-NRBP1, TSC22D2, and TSC22D4 immunoprecipitated from GFP-NRBP1 knock-in HEK293 cells treated with 0.5 M sorbitol for various time points. Whole-cell lysates were also analyzed to assess phosphorylation dynamics. Data represents representative blots of two independent experiments, each with two technical replicates. **(F)** Immunoblot analysis showing the impact of WNK463 (pan-WNK inhibitor) and WNKC11/12 (WNK1/3-specific inhibitors) on NRBP1 phosphorylation and the broader WNK pathway. Both whole-cell lysates and immunoprecipitated NRBP1 complexes were analyzed. Representative blots from one experiment are shown, with two independent experiments performed, each with two technical replicates.

As our NRBP1 phosphospecific pThr232 was not sufficiently sensitive to detect phosphorylated NRBP1 in the whole cell extract, we generated N-terminal tagged GFP-NRBP1 knockin HEK293 cells, to permit facile immunoprecipitation of the endogenous NRBP1 complex using high affinity GFP nanobody, prior to immunoblotting. A time course analysis revealed that following treatment of HEK293 cells with 0.5 M sorbitol a significant increase in phosphorylation of NRBP1 at Thr232 was observed within 1 min that was maintained for 30 min (Fig 4E). We also noted that sorbitol treatment induced a band shift of co-immunoprecipitated TSC22D4 (Fig 4E). Our data suggests that the bandshits of TSC22D4 is a result of phosphorylation, as incubation of immunoprecipitated GFP-NRBP1 with lambda phosphatase reduced this band shift (SFig 4B).

We next found that the kinase domain of WNK1 and WNK3 phosphorylated NRBP1 at Thr232 i*n vitro* (SFig 4C) and that the pan WNK isoform inhibitor, WNK-463 (*65*), as well as two WNK1/WNK3 selective inhibitors (WNK C-11, WNK C-12) (*49*), blocked sorbitol induced phosphorylation of NRBP1 at Thr232 in HEK293 cells (Fig 4F). None of the WNK1 inhibitors impacted the bandshift in TSC22D4 observed with sorbitol, suggesting that this phosphorylation is mediated independently from WNK isoforms (Fig 4F).

### NRBP1 interactome ± sorbitol treatment

To analyze the proteins that associate with endogenous NRBP1 we undertook mass spectrometry analysis of endogenous GFP-NRBP1 immunoprecipitated from HEK293 cells treated with ± 0.5 M sorbitol for 30 min. Cells were lysed in a buffer containing NP-40 (0.5% by vol) and 0.15 M NaCl and following immunoprecipitation of GFP-NRBP1, samples were digested with a mixture of trypsin and LysC protease, processed for quantitative data independent acquisition mass spectrometry and data analyzed through DIA-NN (*66*) (Fig 5A). Control immunoblotting confirmed successful immunoprecipitation of GFP-NRBP1 along with TSC22D4 and that sorbitol induced phosphorylation of SPAK and upshift of TSC22D4 (SFig 5A). Principal component analysis revealed 4 distinct groups mapping with the wild type of control and GFP-NRBP1 cells treated with ± sorbitol (SFig 5B) with significant enrichment of peptides in the GFP-NRBP1 cell lines compared to wild type (SFig 5C). We next analyzed proteins that co-immunoprecipitated specifically with GFP-NRBP1 in non-sorbitol treated cells, with data visualized using the CURTAIN interactive volcano plot software (*67*), in which the data could be viewed and analyzed using the web link provided in the figure legend (Fig 5B). Proteins enriched >4-fold in the GFP-NRBP1 immunoprecipitates from unstimulated cells were, NRBP1, NRBP2, TSC22D1, TSC22D2, TSC22D3, TSC22D4, ELOB, ELOC and a group of proteins involved in the PI3K signaling system, namely IRS4, PIK3R1, PIK3R2, PIK3R3 and not significantly impacted by sorbitol stimulation(Fig 5C). Several other proteins were also enriched to a lower extent in the GFP-NRBP1 immunoprecipitates from unstimulated cells and not affected by sorbitol administration namely, SPAK, MAP7D2, NME3, SPACDR, TBC1D30, LOXL1, MYO5B, OAS3, TUBB4A, SMARCA1, FAM117B, FAM111B and HOXC11 (SFig 6).

**Figure 5:**
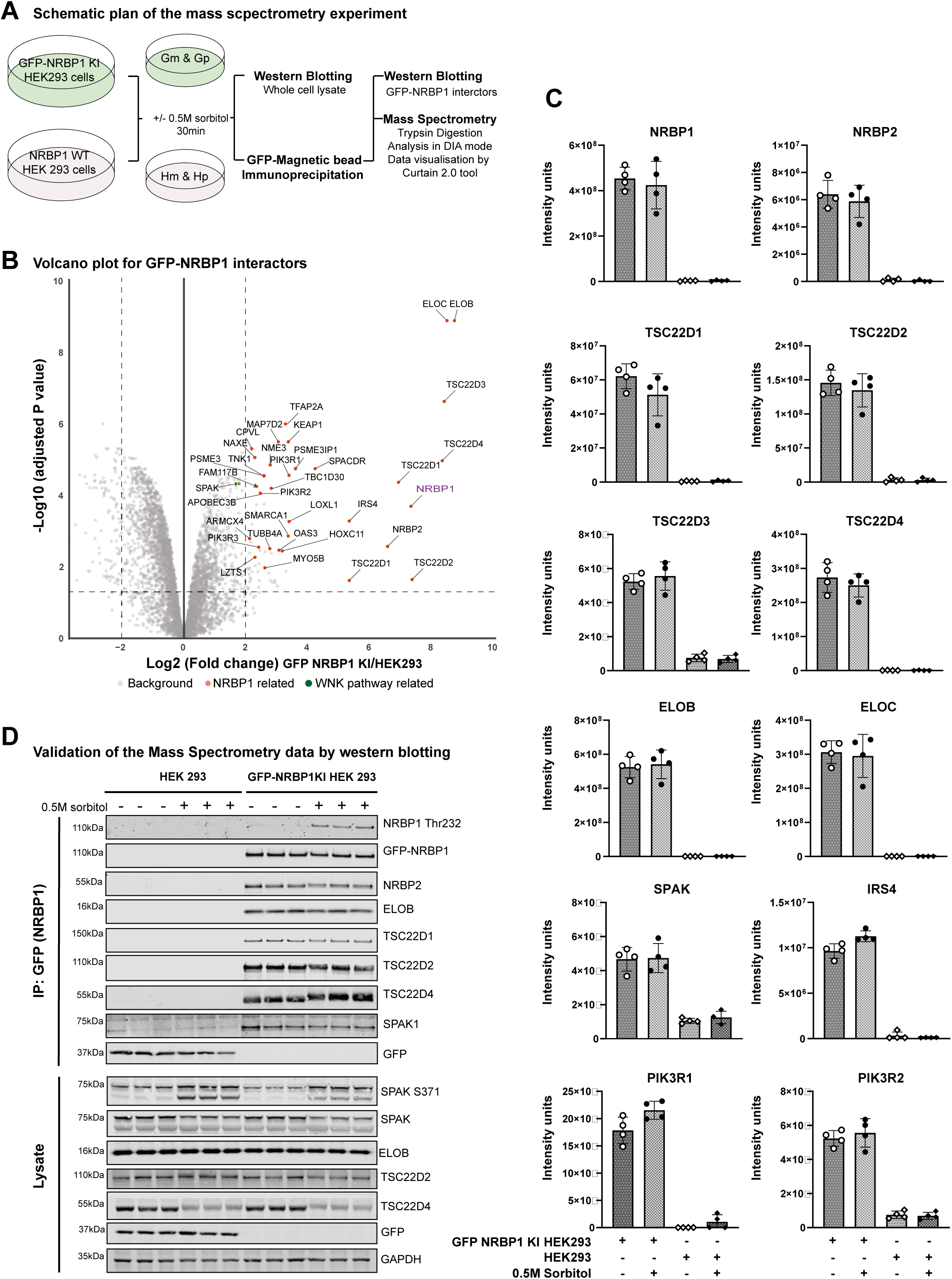
NRBP1 interactors. **(A)** Schematic representation of the mass spectrometry protocol used to identify NRBP1 interactors under basal and sorbitol stress conditions. GFP-NRBP1 knock-in HEK293 cells and wild-type (WT) HEK293 cells (negative control) were treated with or without 0.5 M sorbitol for 30 minutes (N=5 for each group). Following GFP-magnetic bead pulldown and washing, samples were processed for mass spectrometry using S-trap microcolumns according to the manufacturer’s protocol. Peptides were identified in DIA mode, analyzed in Python, and visualized with the Curtain tool. Note: Gm-GFP NRBP1 cells without sorbitol, Gp-GFP NRBP1 cells with sorbitol, Hm-WT NRBP1 cells without sorbitol and Hp-WT NRBP1 cells with sorbitol. **(B)** Volcano plot showing proteins significantly enriched (≥4-fold) as NRBP1 interactors under unstimulated conditions. GFP-NRBP1 intensities were normalized to WT-NRBP1, and statistical significance was determined using Benjamini-Hochberg correction. Proteins with corrected P-values <0.05 were considered significant. Data from four technical replicates per group were visualized using the Curtain 2.0 tool. Curtain link -https://curtain.proteo.info//a38745d4-927f-411e-a89d-b10e6bdd2f9c. **(C)** Box plots showing normalized protein intensities of the interactors identified as significant in the volcano plot (B). **(D)** Selected NRBP1 interactors identified in MS were validated by western blotting. Experiments were repeated twice with three technical replicates each, confirming the enrichment of interactors detected in the mass spectrometry analysis.

**Figure 6:**
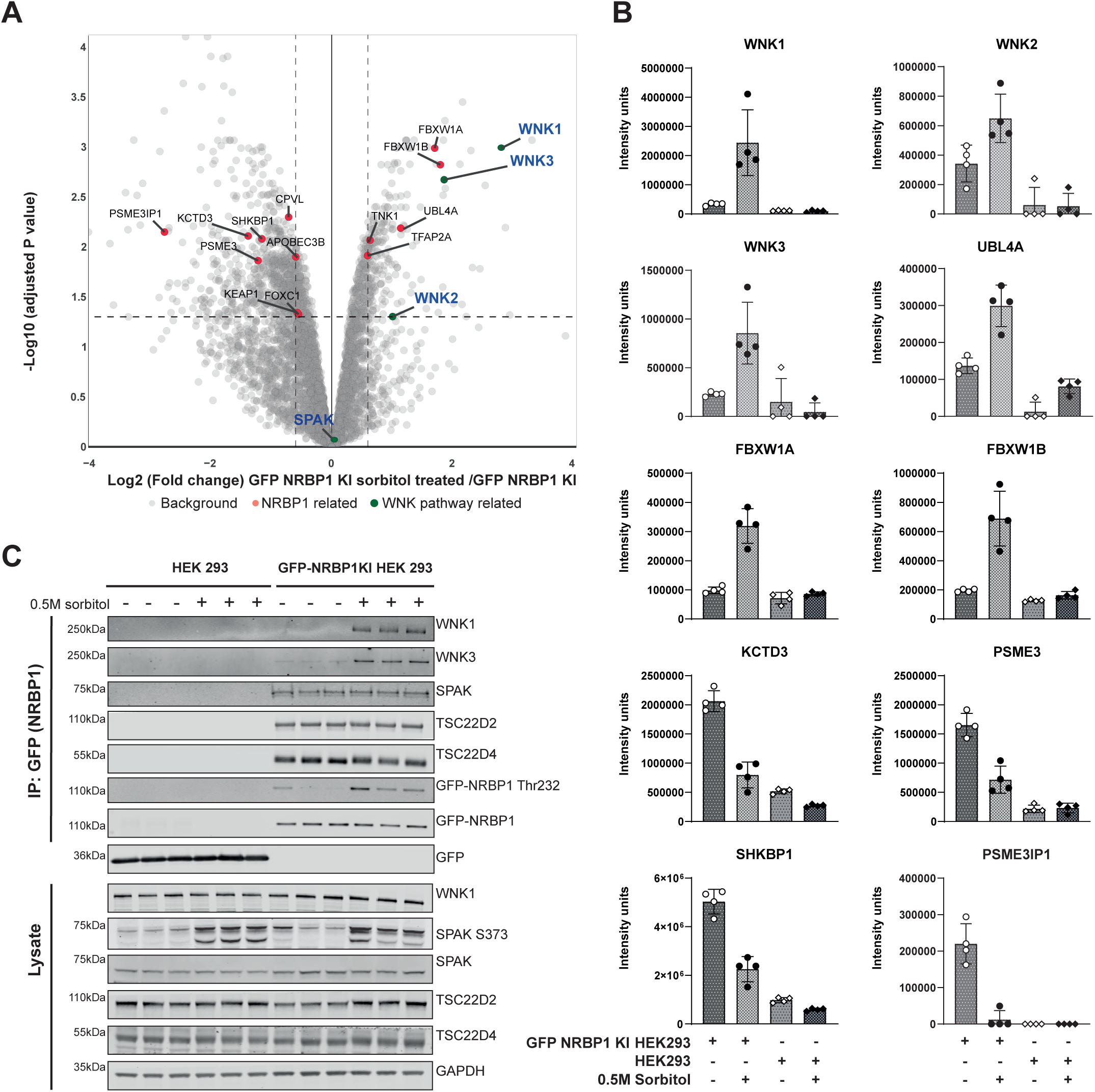
NRBP1 differential interactors. **(A)** Volcano plot displaying NRBP1 interactors significantly enriched (≥2-fold) following treatment with 0.5 M sorbitol. GFP-NRBP1 intensities were normalized to WT-NRBP1 within the sorbitol-treated group. Statistical significance was determined using Benjamini-Hochberg correction, with proteins considered significant at a corrected P-value <0.05. Data represents four technical replicates per group, visualized using the Curtain tool. Curtain-link https://curtain.proteqo.info//6a8a5480-b5db-4c1e-ab94-c7403a37f850 **(B)** Box plots showing normalized protein intensities for the significant hits identified in the volcano plot (A). **(C)** Key NRBP1 differential interactors identified by mass spectrometry were validated by western blotting. Experiments were performed twice, each with three technical replicates.

Consistent with our proximity ligation data revealing that sorbitol induces the association of WNK1 and NRBP1 (Fig 1D,E), we observed that sorbitol stimulation induced significant association of NRBP1 with WNK1 (7-fold) as well as WNK2 (2-fold) and WNK3 (3.5-fold) in addition to 3 proteins involved in ubiquitin pathways UBL4A (2-fold), βTrCP1/FBXW1A (3.5-fold) and βTrCP2/FBXW1B (3.5-fold) (Figure 6A&B). Immunoblotting confirmed that sorbitol induced association of NRBP1 with WNK1 and WNK3, (Fig 6C). We also observed that several proteins (including PSME3, PSME3IP1, SHKBP1 and KCTD3) interaction with GFP-NRBP1 was reduced following sorbitol stimulation (Figure 6A&B).

### Rapid degradation or knock-out of NRBP1 inhibits WNK activation and phosphorylation of SPAK/OXSR1 and NKCC1

To define the role that NRBP1 plays in regulating the WNK pathway in cells, we first employed a targeted protein degradation approach to rapidly degrade NRBP1. We used a CRISPR knock-in approach to attach a BromoTag, an inducible degron motif (*68*), to the N-terminus of NRBP1 in HEK293 cells. Employing this approach, the AGB1 heterobifunctional degrader compound, hijacks the VHL E3 ligase to induce rapid ubiquitylation and subsequent degradation of the BromoTag-NRBP1 protein (*68*) (Fig 7A). We observed that doses of 100 or 300 nM AGB1 for 3h, reduced BromoTag-NRBP1 levels > 95% without impacting NRBP2 levels (Fig 7B, SFig 7A). Although these cells were not stimulated with sorbitol, knockdown of NRBP1 was accompanied by a significant reduction in levels of basal T-loop phosphorylation of SPAK/OXSR1 (Fig 7B, SFig7B). As a control we used 300 nM of the inactive cis-AGB1 degrader control compound (*68*) that induced no degradation of BromoTag-NRBP1 and also had no effect on the T-loop phosphorylation of SPAK/OXSR1 levels (Fig 7B). We next treated BromoTag-NRBP1 HEK293 cells with 100 nM AGB1 for between 0.5 to 4 h followed by 0.5 M sorbitol for 30 min, in the presence of AGB1 (Fig 7C). Under these conditions, the BromoTag-NRBP1 protein levels were reduced by ∼80% within 30 min and > 95% within 1h (Fig 7C). Reduction in BromoTag-NRBP1 levels were also accompanied by a corresponding decrease in the sorbitol induced T-loop phosphorylation of SPAK/OXSR1, as well as phosphorylation of the SPAK/OXSR1 substrate NKCC1 (Thr212/Thr217) (Fig 7C). To confirm these findings, we also generated 6 independent clones of NRBP1 knock-out HEK293 cells and observed that all these cell lines displayed significantly reduced levels of phosphorylation of SPAK/OXSR1 following sorbitol stimulation (SFig 7C).

**Figure 7:**
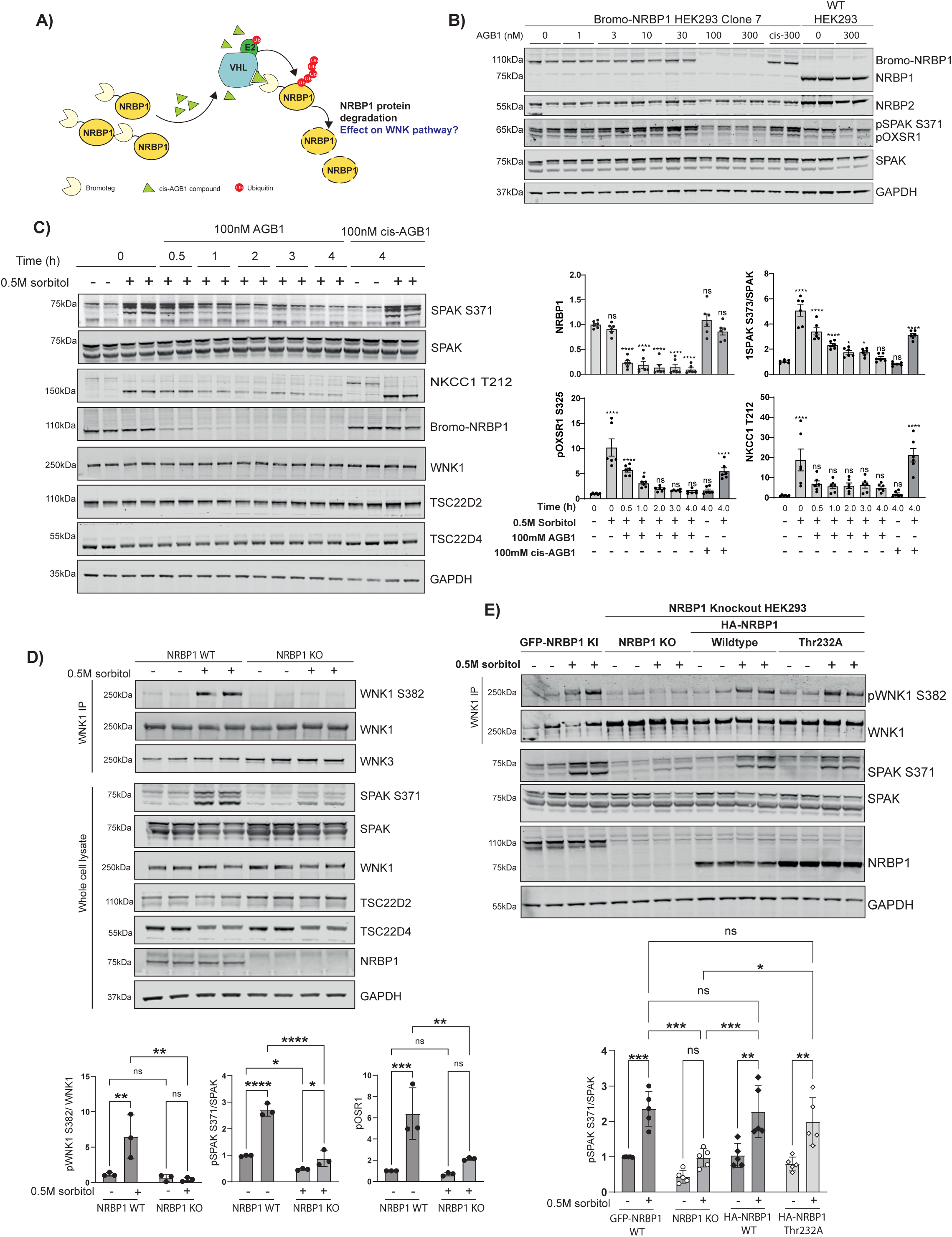
NRP1 regulates the WNK1 pathway. **(A)** Cartoon illustration depicting the inducible degradation system used for NRBP1. NRBP1 was N-terminally tagged with the bdTag (BromoTag), which, upon binding to the AGB1 compound, is directed to the proteasome for degradation. **(B)** Bromotag-NRBP1 knock-in (KI) HEK293 cells were treated with increasing concentrations of AGB1 (active compound) or cis-AGB1 (negative control inactive compound) for 3 h. Lysates were subjected to immunoblotting to assess dose-dependent degradation of NRBP1. **(C) Left Panel**: Time-dependent degradation of NRBP1 was analyzed alongside the WNK1 signaling pathway. Bromotag-NRBP1 KI HEK293 cells were treated with 100 µM AGB1, followed by 0.5 M sorbitol for 30 minutes. The impact on WNK pathway markers was assessed by immunoblotting. **Right Panel**: Densitometric analysis of immunoblot results showing the changes in WNK pathway activation. Data represent three independent experiments, each with two technical replicates.**(D) (Upper)**The impact of NRBP1 knockout on WNK1 activation (pWNK1-S382) was assessed by immunoprecipitating endogenous WNK1 and immunoblotting.**(Lower)** Densitometric analysis of the western blots were performed**. (E)** The impact of NRBP1 knockout and rescue with either WT NRBP1 or the Thr232A mutant was assessed on the WNK pathway by Immunoblots analysis. Statistical analysis was performed using two-way ANOVA with Sidak’s multiple comparison test (**P < 0.01, ***P < 0.001, ****P < 0.0001).

We also studied the impact of NRBP1 knock-out on the sorbitol induced phosphorylation of the T-loop Ser382 site of WNK1. This was achieved by immunoprecipitating WNK1 from wild type and NRBP1 knock-out cells treated with ± 0.5 M sorbitol for 30 min and immunoblotting with a phosphospecific pSer382 antibody. The results strikingly revealed that loss of NRBP1 markedly reduced sorbitol induced WNK1 Ser382 phosphorylation (Fig 7D).

We next stably re-expressed wild type NRBP1 or NRBP1[Thr232A] into NRBP1 knock-out cells, using a retrovirus system. Expression of wild type NRBP1 in the NRBP1 knock-out cells restored WNK-mediated T-loop phosphorylation of SPAK/OXSR1 in sorbitol treated cells (Fig 7E). Similar rescue of SPAK/OXSR1 phosphorylation was observed following expression of the NRBP1[Thr232A], indicating that the impact NRBP1 has on SPAK/OXSR1 T-loop phosphorylation is independent of NRBP1 Thr232 phosphorylation (Fig 7E).

### Recombinant NRBP1 activates WNK4

We next investigated whether recombinant NRBP1 expressed in *E. coli* could activate WNK isoforms. For these experiments we employed a fragment of WNK4 encompassing the kinase domain (residues 1 to 449) that was expressed in insect cells, as unlike WNK1, WNK2 and WNK3 isoforms expressed in bacteria, this fragment of WNK4 was not constitutively phosphorylated at its T-loop site (Ser335, equivalent to Ser382 in WNK1) (*38*). We combined NRBP1 (full length 2 μM), in the presence or absence of WNK4 (residues 1-449, 500 nM) and kinase-inactive OXSR1[D16A] (full length, 2 μM). The mixtures were incubated for 40 min in the presence of Mg-ATP at 30 °C. Reactions were stopped with SDS sample buffer and phosphorylation of NRBP1 at Thr232, WNK4 at Ser335 and OXSR1 at Ser325 and Thr185 monitored by immunoblotting using phosphospecific antibodies. Consistent with insect cell expressed WNK4 being devoid of activity, in the absence of NRBP1, it only phosphorylated kinase inactive OXSR1 at its T-loop residue to a low level (Fig 8). Furthermore, as expected, the NRBP1 pseudokinase did not phosphorylate kinase-inactive OXSR1 in the absence of WNK4 (Fig 8). However, by adding NRBP1 to WNK4 and kinase inactive OXSR1, we observed a marked phosphorylation of WNK4, NRBP1 and OXSR1 T-loop residues (Fig 8). Our parallel study also provides further evidence in a cellular model as well as in mice that NRBP1 is capable of activating WNK4 kinase in kidney cells of relevance to controlling human blood pressure (*51*).

**Figure 8:**
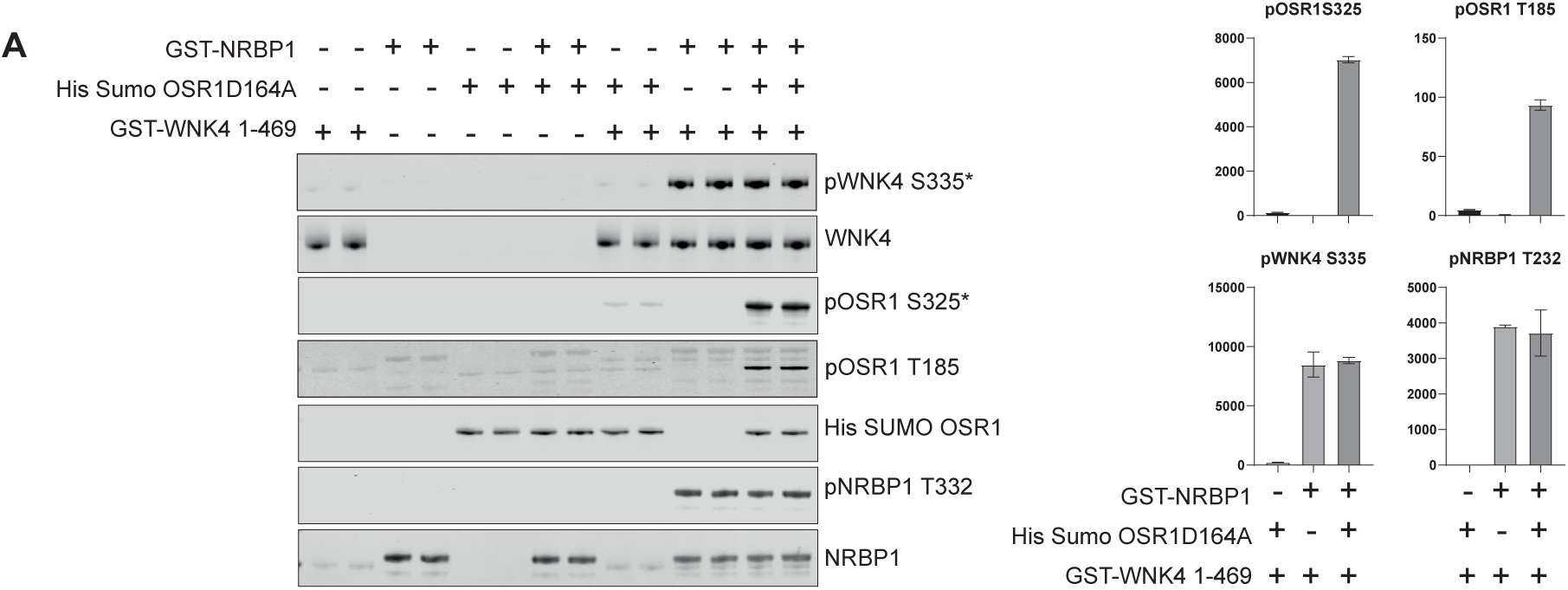
WNK4 is activated *in vitro* by NRBP1. The activation of WNK4 by NRBP1 was assessed through an *in vitro* kinase assay. The reaction included GST-WNK4 (residues 1–469), GST-NRBP1, and His-Sumo-OXSR1 (D164A, a catalytically inactive mutant). The reaction products were analyzed by immunoblotting to detect WNK4 activation as well as phosphorylation of OXSR1 and NRBP1. Representative results from two independent experiments, each performed with two technical replicates, are shown.* indicates that the blotting was performed with the pWNK1-S382 and pSPAK-S371 antibody which detects the consensus phosphosite in WNK4 and OXSR1, respectively.

### AlphaFold-3 analysis of the NRBP1:TSC22D4: WNK1:SPAK complex

Next, we undertook AF3 modelling of the WNK-NRBP1 complex. The AF3 model of NRBP1 suggests that it forms a homodimer via its pseudokinase domain (SFig 8A) or a heterodimer with the kinase domain of WNK1 (SFig 8B). In the AF3 model of full length WNK1, most of the non-catalytic residues are assigned as unstructured apart from the two CCT-like domains (CCTL1 and CCTL2) (SFig 8C). In contrast, in the AF3 model of the heterotrimeric NRBP1:WNK1: SPAK complex, the non-catalytic residues of WNK1 are largely ordered (SFig 8D). In this structure the RΦ-motif motif-2 of WNK1 (residues 1257-1260, RFIV) interacts with the SPAK-CCT domain (SFig 8E), in agreement with earlier experimental work (*4*). In the NRBP1:WNK1: SPAK structure the WNK1-CCTL1, WNK1-CCTL2 and NRBP1-CCT domains are all unoccupied (SFig 8E). Finally, we modeled a complex of NRBP1:WNK1: SPAK with homodimeric TSC22D4, which results in an ordered TSC22D4 dimer in which the TSC22D4 RΦ-motif-A forms an interaction with the WNK1-CCTL1, and the RΦ-motif-B interacts with the NRBP1-CCT domain (Fig 9A, 9B). In the NRBP1:WNK1:SPAK: homodimeric TSC22D4 complex, the WNK1 bound ATP gamma-phosphate group is poised towards three residues on SPAK (Thr231, Ser371 and Ser385), that comprise the 3 reported WNK1 phosphorylation sites (*2*) (Fig 9C).

**Figure 9:**
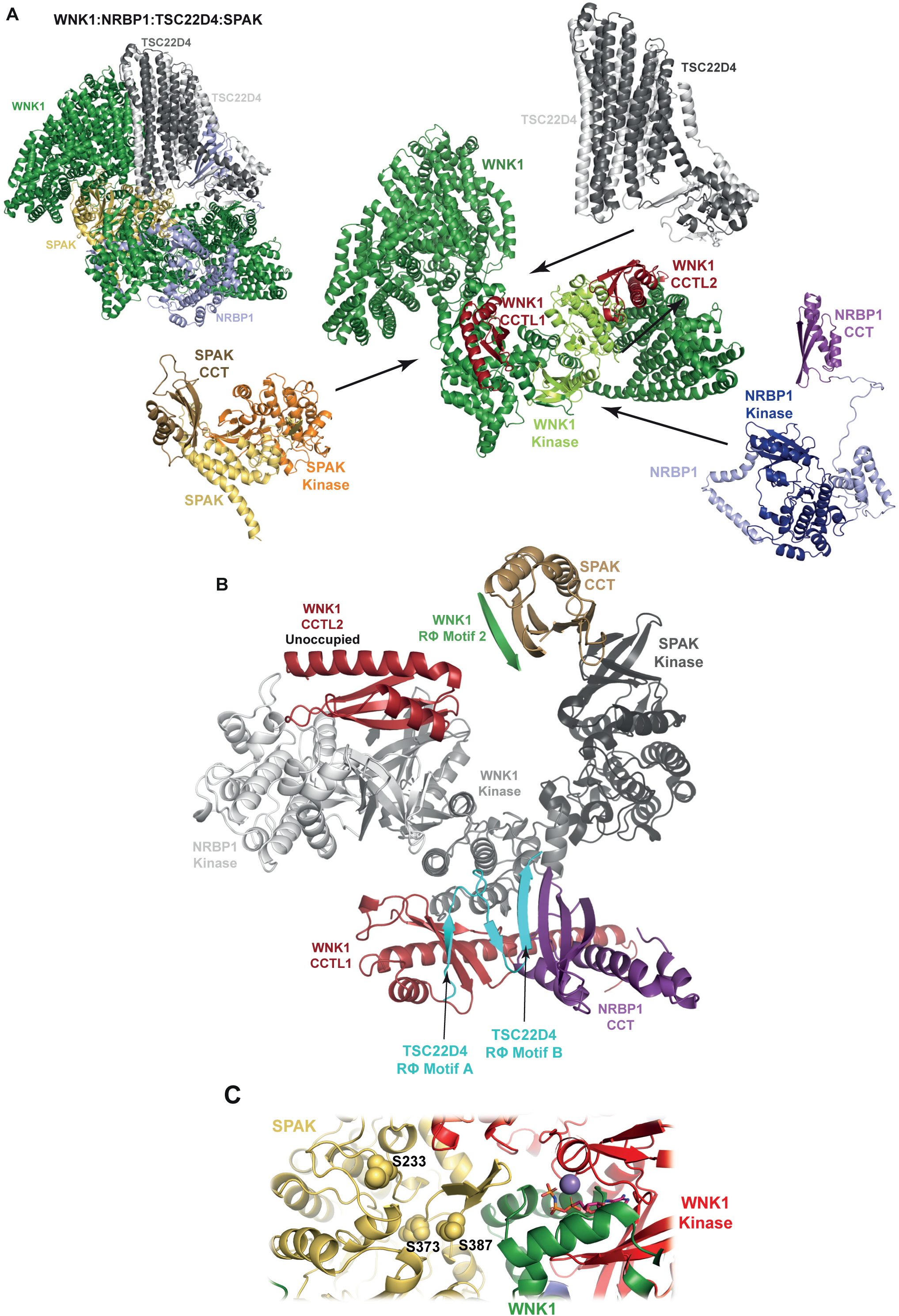
AlphaFold3 modelling of the WNK1-SPAK NRBP1 & TSC22D4 complex. **(A)** A structural model of WNK1 in complex with SPAK, NRBP1, and a TSC22D4 homodimer, generated using AlphaFold3. The model is shown both as a complete assembly and disassembly into individual components, highlighting key structural domains. The conserved CCT domains in NRBP1 and TSC22D4 are labeled. **(B)** As in (A), but showing only the kinase domains, CCT domains, and RΦ motifs within the WNK1-NRBP1-TSC22D4 complex. **(C)** Detailed view of the active site of WNK1 kinase, showing ATP positioned near key phosphorylation sites on SPAK (Thr231, Ser371, and Ser385).

## Discussion

NRBP1 was first cloned in 2000 as a protein possessing a “kinase-like” motif and was predicted to bind to nuclear receptors due to the presence of LXXLL motifs at its C-terminus (*59*), which to our knowledge has not been demonstrated experimentally. Subsequent work reported various roles for NRBP1 including interacting with Rac3 GTPase (*69*), controlling eye development (*70*) and operating as a negative regulator of gene transcription and tumor progression (*71*). NRBP1 is ubiquitously expressed and knockout of NRBP1 in mice results in early embryonic lethality at day E7.5 (*48*). A genetic analysis in Caenorhabditis elegans as well mice, suggested that NRBP1 influenced proliferation and homeostasis of intestinal progenitor cells and tumor formation (*48*). This study also reported for the first time that NRBP1 co-immunoprecipitated with TSC22D2 and TSC22D4 as well as key components of the ubiquitination machinery including Elongin B and Elongin C (*48*) which form a complex that operates as a substrate adaptor for the Cullin-2 and Cullin-5 E3 ligases (*72*). Our data confirm that Elongin B and Elongin C are constitutively associated with NRBP1 in a manner that is not impacted by sorbitol (Fig 5). We found that sorbitol promotes interaction of NRBP1 with WNK1 as well as other ubiquitin pathway components including βTrCP1/FBXW1A and βTrCP2/FBXW1B that function as members of the F-box substrate adaptors of Cullin-1 E3 ligase complex that operates independent of Elongin B and Elongin C (*72*). Another study has suggested that NRBP1 Cullin containing complexes can regulate the ubiquitylation of two transmembrane proteins termed BRI2 and BRI3 that control the activity of α- and β-secretase (*49*). Given the association of NRBP1 with so many ubiquitin pathway components, in future work it would be important to explore whether NRBP1 could impact ubiquitylation of WNK pathway components. Our current data indicates that knock-out or conditional knock-down of NRBP1 does not impact expression levels of WNK1, SPAK or OXSR1 proteins, as assessed by immunoblotting.

NRBP1 is classified as a class 1 pseudokinase as it lacks 3 key motifs (DFG, VAIK and HRD) present on all protein kinases that are required for catalysis of protein phosphorylation. NRBP1 was also shown to lack catalytic activity (*53*) and is incapable of binding ATP or cations (*73*). AF3 modeling suggests that the pseudokinase domain of NRBP1 can homodimerize, or heterodimerize with the kinase domain of WNK1 (SFig 9). Due to the large interface of these dimer interactions, it would be hard to suppress these interactions by simple mutations. Proximity ligation data and co-immunoprecipitation analysis (Fig 1 & Fig 4) reveals that osmotic stress induces a rapid and robust interaction of NRBP1 with WNK1 (Fig 1&6) which has also been reported in a recent study (*50*). Further work is required to understand the mechanism by which sorbitol triggers the association of WNK1 and NRBP1, which has been suggested to be driven by intrinsically disordered regions within WNK1 and the TSC22D proteins (*50*). Our data also indicates that sorbitol induces the phosphorylation of TSC22D4 causing an upward electrophoretic mobility shift, which is independent of the WNK kinase activity as it is not impacted by WNK inhibitors (Fig 4). Future work is also required to map these phosphorylation sites, identify the upstream kinases and whether they play any role in controlling the interaction of NRBP1 with WNK isoforms.

Our results indicate that one of the consequences of NRBP1 association with WNK isoforms is phosphorylation of Thr232 residue in the T-loop of NRBP1 that becomes rapidly phosphorylated in a manner that is blocked by a pan WNK kinase inhibitor as well as a WNK1/WNK3 dual inhibitor (Fig 4). Our data indicate that association of NRBP1 to WNK1 occurs rapidly, within 1 min of osmotic stress, and presumably this binding is required for WNK1 to phosphorylate NRBP1 at Thr232. Monitoring of Thr232 phosphorylation could be used as a biomarker for the association of WNK1 with NRBP1. Our finding that mutation of Thr232 residue to Ala does not impact the ability of NRBP1 to rescue SPAK/OXSR1 phosphorylation when reconstituted into NRBP1 knock-out cells, suggests that Thr232 phosphorylation is not required for NRBP1 to activate the WNK1 pathway.

Our data, point towards NRBP1 controlling the activation of the basal WNK pathway as well as its sorbitol stimulated activity as both rapid knockdown or knock-out of NRBP1 substantially suppressed basal and sorbitol induced WNK autophosphorylation and the WNK mediated phosphorylation and activation of SPAK/OXSR1. The loss of WNK pathway activity can be rescued by expression of NRBP1 in knock-out HEK293 cells. Our finding that T-loop phosphorylation and activation of a bacterially expressed inactive fragment of WNK4 can be stimulated by NRBP1 in the presence of OXSR1 (Fig 8), is consistent with NRBP1 functioning as an upstream activator of the pathway. Our recent studies in which NRBP1 was knocked-out in distal convoluted tubule cells of the kidney, also support that this inhibits the WNK4 signaling pathway’s ability to regulate the level and activity of the thiazide sensitive NCC ion cotransporter (*51*). Our data suggests that NRBP1, WNK1 and SPAK/OXSR1 together with TSC22D2/4, could form a complex in which the conformation of the WNK1 kinase is stabilized in the active conformation, capable of autophosphorylating Ser382 in the T-loop residue, a prerequisite to activate WNK isoforms. This data is also consistent with our proximity ligation (Fig 1) mass spectrometry (Fig 5&6) and immunoprecipitation data showing that in cells treated with sorbitol TSC22D2/4-NRBP1-WNK1-SPAK/OXSR1 could form a complex.

The TSC22D2/4:NRBP1:WNK1:SPAK complex possesses four CCT domains (NRBP1-CCT, WNK1-CCTL1, WNK1-CCTL-2 and SPAK-CCT). AF3 modeling of the complex indicates that three out of the four CCT domains are occupied by binding to specific RΦ-motifs with only the WNK1 CCTL2 domain remaining unoccupied. The SPAK-CCT domain is complexed to the WNK1 RΦ-motif-2 (residues 1257-1260, RFIV, Fig 2B) consistent with previous experimental work (*4*). The NRBP1 CCT domain is complexed to the RΦ-motif-B of TSC22D4, whereas the RΦ-motif-A of TSC22D4 is complexed to the WNK1 CCTL1. These CCT-RΦ-motif interactions likely provide important structural integrity to the TSC22D2/4:NRBP1:WNK1:SPAK complex. The finding that the gamma phosphate moiety of ATP within the WNK1 kinase domain on the TSC22D2/4:NRBP1:WNK1:SPAK complex, is oriented towards the three well characterized SPAK phosphorylation sites (Thr231, Ser371 and Ser385), located on two distinct regions of SPAK, provides further evidence to support the reliability of the model. It would be important in future work to undertake experimental structural analysis to better understand the regulation and function of the TSC22D4:NRBP1:WNK1:SPAK complex. How the NRBP1 pseudokinase controls the conformation and activity of WNK kinases is not defined by the AF3 modeling we have undertaken.

There are two other TSC22D isoforms family members namely TSC22D1 and TSC22D3 that were picked up in our WNK1 proximity ligation or NRBP1 immunoprecipitation studies. TSC22D1 possesses the same conserved RΦ-motif-A (RFRV) and RΦ-motif-B (RWTC) present in TSC22D2 and TSC22D4. However, neither of these motifs are present in TSC22D3 but it could form heterodimers with other TSC22D isoforms. A recent study has suggested that TSC22D1 interacts with the CCT domain of NRBP1 (*61*). We have also observed that immunoprecipitation of NRBP1 results in the co-immunoprecipitation of NRBP2, a closely related pseudokinase isoform. It is possible that NRBP1 and NRBP2 could heterodimerize. In future, it would be important to further explore whether NRBP2 works together with NRBP1 to activate the WNK pathway or whether these pseudokinases play distinct roles. We were unable to generate NRBP1 and NRBP2 double knock-out HEK293 cells, suggesting that these proteins could play an essential redundant role.

There are several well-characterized examples of pseudokinases interacting with and activating protein kinases. Notable cases include the LKB1 tumor suppressor kinase, which activates AMPK family members and is regulated by the pseudokinases STRADα and STRADβ ((*74*)). The ability of STRAD isoforms to activate LKB1 is further enhanced by the adaptor protein MO25 ((*75*)). Another example is ErbB3, which allosterically activates other EGFR receptors ((*76*)). In JAKs, the JH2 pseudokinase domain modulates the conformation and activity of the adjacent JH1 protein kinase domain ((*77*)). Similarly, the KSR1/2 pseudokinases bind and regulate the activity of Raf and MEK isoforms ((*78*)). In each of these cases, the pseudokinase adopts a conformation that mimics the active state of a catalytically competent kinase in response to specific stimuli. This conformational shift enables the pseudokinase to interact with and allosterically activate its associated kinase. NRBP1, and likely NRBP2, working in conjunction with WNK kinases, represent additional examples of this regulatory pseudokinase-kinase signaling module. In future research, it will be crucial to explore whether osmotic stress, possibly through the formation of biomolecular condensates or another mechanism, could induce the NRBP1 pseudokinase to adopt an active conformation capable of interacting with and activating WNK isoforms. Such insights would significantly advance our understanding of how osmotic stress is sensed and how it triggers the activation of the WNK signaling pathway.

In summary, our data suggests that NRBP1 and TSC22D2/4 form a constitutive complex that interacts with WNK isoforms as well as SPAK/OXSR1 in response to osmotic stress. This interaction appears to promote the activation of WNK isoforms, which in turn phosphorylate and activate SPAK/OXSR1, while also phosphorylating NRBP1 at Thr232. The functional significance of NRBP1 phosphorylation at Thr232 remains unclear, but it serves as a useful marker for the association of NRBP1 with WNK isoforms. Further research is needed to elucidate how osmotic stress is sensed, how it links to biomolecular condensates, and how it regulates the interaction between NRBP1 and WNK isoforms.

## MATERIALS AND METHODS

### Plasmids

Standard recombinant DNA techniques were employed as previously described. Constructs for transient transfections were subcloned into pcDNA5-FRT/TO or pCMV5 vectors. For retroviral transfections, the pBABE puro vector was utilized. All constructs are available for request from the Medical Research Council (MRC) Phosphorylation and Ubiquitylation Unit (PPU) Reagents webpage at MRC PPU Reagents (https://www.ppu.mrc.ac.uk/). The unique identifier (DU) numbers listed below provide direct access to the cloning strategies and sequence information.

### Antibodies

#### Primary Antibodies for Western Blotting

Anti-phospho-NRBP1 Thr232: DA079 (MRC PPU Reagents, sheep polyclonal, 1 μg/ml; with 10 μg/ml non-phospho-peptide). Anti-TSC22D2: DA056 (MRC PPU Reagents, sheep polyclonal, 1 μg/ml). Anti-TSC22D1: 101501-T36 (Sinobiologicals, rabbit polyclonal, 1:2000). Anti-phospho-WNK1 Ser382: S099B (MRC PPU Reagents, sheep polyclonal, 1 μg/ml; with 10 μg/ml non-phospho-peptide). Anti-WNK1: Abcam 37687, 1:2000. Anti-phospho-SPAK Ser371: S670B (MRC PPU Reagents, sheep polyclonal, 1 μg/ml; with 10 μg/ml non-phospho-peptide). Anti-SPAK: S637B (MRC PPU Reagents, sheep polyclonal, 1 μg/ml). Anti-phospho-NCC Thr-53: PA5-95674 (Thermo Fisher, rabbit polyclonal, 1:1000; can cross-react with NKCC1 Thr212). Anti-NRBP1: SAB1408678 (Sigma-Aldrich, rabbit polyclonal, 1:1000). Anti-NRBP2: DA240 (MRC PPU Reagents, sheep polyclonal, 1 μg/ml; with 10 μg/ml non-phospho-peptide). Anti-WNK4: NB600-284 (Novus Biologicals, rabbit polyclonal, 1:2000). Anti-HA: Roche 3F10 (rat monoclonal, 1:3000). Anti-FLAG: (mouse monoclonal, 1:3000). Anti-GFP: 3H9 (Chromotek, rat monoclonal, 1:3000). GAPDH: Sc-32233 (mouse polyclonal, 1:5000). α-Tubulin: 3873 (Cell Signaling Technology, mouse monoclonal, 1:10,000).

#### Peptides for Sheep Phosphospecific Antibodies

Sourced from Peptides & Elephants.

#### Antibody Production

Sheep polyclonal antibodies were produced by immunizing sheep with peptide antigens (Peptides & Elephantas). These peptides were conjugated to carrier proteins, keyhole limpet hemocyanin (KLH) and bovine serum albumin (BSA), through C-terminal linkage with 6-aminohexanoic acid (Ahx), as outlined in Harlow and Lane (1988) Antibodies: A Laboratory Manual (Cold Spring Harbor Laboratory Publications). Sheep were immunized with the antigen and subsequently received up to four booster injections, spaced 28 days apart. Blood samples were collected seven days after each injection. Antibodies were affinity-purified from the sera using the same phosphopeptides used for immunization. Briefly, serum was heat-inactivated at 56°C for 20 min and then filtered through a 0.45 µm filter. The inactivated serum was diluted 1:1 with 50 mM Tris/HCl pH 7.5 containing 2% Triton X-100. The diluted serum was passed through a column containing the antigen-conjugated resin, with the flowthrough collected twice and stored at -20°C as a backup. The resin was washed with Peptide Wash Buffer (50 mM Tris/HCl pH 7.5, 0.5 M NaCl) until the absorbance at 595 nm (OD595) was less than 0.003 to eliminate non-specific proteins and antibodies. Antibodies were eluted in 1 ml fractions using 50 mM Glycine pH 2.5 into 1.5 ml Eppendorf tubes containing 200 µl of 1 M Tris/HCl pH 8.0 to neutralize the elution buffer. The antibody concentration was determined using the Bradford assay and then dialyzed against 1X PBS overnight. The final concentration was adjusted to above 0.1 mg/ml, and aliquots of the purified antibodies were stored at –20°C.

#### Cell Culture, Transient Transfection, and Cell Lysis

HEK-293 human embryonic kidney cells (ATCC CRL-1573) were cultured in Dulbecco’s Modified Eagle’s Medium (DMEM; GIBCO 11960–085) supplemented with 10% fetal bovine serum (FBS; Sigma F7524), 2 mM L-glutamine (GIBCO 25030024), and 100 U/mL penicillin-streptomycin (GIBCO 15140122). Cells were maintained at 37°C in a humidified atmosphere with 5% CO2 and regularly tested for mycoplasma contamination. For transient transfections, 10 μl of polyethylenimine (PEI, 1 mg/ml; Polysciences Inc 24765) was mixed with 2 μg of plasmid DNA in 0.5 mL of Opti-MEM (GIBCO 31985–062). The mixture was vortexed gently for 20 seconds and incubated at room temperature for 30 min. After incubation, the transfection mixture was added dropwise to the cell culture medium in a 60 cm² dish. Cells were treated and lysed 24-36 h post-transfection. Following treatment, cells were lysed directly on the plate using ice-cold lysis/immunoprecipitation buffer (50 mM Tris-HCl pH 7.5, 1% (by vol) NP-40, 100 mM NaCl, 1 mM EDTA, 1 mM sodium orthovanadate, 50 mM sodium fluoride, 5 mM sodium pyrophosphate, 10 mM sodium β-glycerophosphate, and cOmplete Mini EDTA-free protease inhibitor [Merck 11836170001]) without a prior PBS wash. The same lysis buffer was used for immunoprecipitation. Protein lysates were clarified by centrifugation at 17,500 rpm for 15 min at 4°C, and protein concentration was measured using the Bradford assay.

#### Quantitative Immunoblotting Analysis

For immunoblotting, approximately 20-25 µg of protein lysates and 25% of immunoprecipitated fractions were resolved by electrophoresis on precast NuPAGE 4–12% Bis–Tris Midi Gels (Thermo Fisher) at 120 V using NuPAGE MOPS-SDS running buffer (Thermo Fisher Scientific). Proteins were transferred to nitrocellulose membranes (GE Healthcare, Amersham Protran Supported 0.45 µm) for 90 min at 90 V using a transfer buffer (48 mM Tris, 39 mM glycine, 20% (by vol) methanol). The membranes were blocked with 5% non-fat milk in 0.1% TBS-T for 1 h at room temperature. Primary antibody incubation was performed either overnight at 4°C or for 1 h at room temperature. Following primary antibody incubation, membranes were washed three times for 10 min each with 0.1% TBS-T. Secondary antibody incubation was performed for 1 h with anti-sheep, anti-rabbit, or anti-mouse IgG secondary antibodies, which were fluorescently labeled with IR680 or IR800.

#### Immunoprecipitation Assay

Immunoprecipitations were conducted as described previously (*17*). Protein lysates (1-4 mg, depending on the experiment) were used for immunoprecipitation with NHS-activated GFP and HA-Sepharose nanobeads from MRC-PPU Reagents and Services. Approximately 20% of the reaction was reserved as input or whole cell lysate. For the immunoprecipitation, 20 μL of 1X PBS-washed bead slurry per 2 mg of lysate was used. The incubation was performed at 4°C for 2 h with gentle agitation at 20 rpm. The beads were then washed sequentially with 1 mL of IP buffer, followed by two washes with ice-cold 1X PBS containing 0.02% (by vol) NP-40. Each wash was followed by centrifugation at 1g for 3 min. Proteins were eluted from the beads by boiling in 50 µL of 2X NuPAGE LDS sample buffer at 60°C for 5 min. The eluate was then filtered using a Spin-X Centrifuge Tube Filter (Costar 8161), and denatured with 1% (by vol) β-mercaptoethanol at 95°C for 5 min.

#### Statistical Analysis

Statistical analyses were conducted using GraphPad Prism (RRID:SCR_002798, version 9.3.1; http://graphpad.com/). The following tests were used: Two-tailed unpaired t-test: Applied for comparisons between two independent groups. One-way ANOVA: Used for comparisons among three or more groups. Post hoc tests were employed where appropriate to determine specific group differences following ANOVA.

### Protein Expression

#### Vectors and Cells

Transform pGEX6P-1 vectors expressing N-terminal Glutathione-S-transferase (GST) tagged proteins (WNK1 (1-661), DU43603; WNK2 (1-627), DU49486; OXSR1 (1-527); NRBP1 (1-535), DU68346; NRBP1 (1-535) T232A, DU68840) into BL21 Codon Plus cells. For GST-WNK3 (1-579), DU4631, use autoinduction media. Culture and Induction: Inoculate a 16-h starter culture into 1 L of Luria-Bertani (LB) broth with 100 µg/mL carbenicillin. Grow at 37°C with shaking at 200 rpm until OD600 reaches 0.8. For GST-WNK3, grow until OD600 reaches 2. Induce protein expression by adding 0.05 mM Isopropyl β-D-1-thiogalactopyranoside (IPTG) and incubate at 18°C with shaking at 200 rpm for 18 h (24 h for GST-WNK3).

#### Cell Lysis and Protein Purification

Pellet cells at 4200 g for 30 min. Resuspend the pellet in 20 mL of ice-cold lysis buffer (50 mM Tris-HCl pH 7.5, 250 mM NaCl, 1% Triton X-100, 1 mM EDTA, 1 mM EGTA, 0.1% β-mercaptoethanol, 0.2 mM PMSF, 1 mM benzamidine). Lyse cells by brief freeze-thaw cycles at -80°C and sonicate (Branson Digital Sonifier) with ten 15-second pulses at 45% amplitude.

#### Clarification and Affinity Purification

Centrifuge the lysate at 35,000 g for 30 min. Incubate the supernatant with 2 mL of pre-equilibrated GSH-Agarose (Abcam) beads at 4°C for 1 h. Wash the beads thrice with a wash buffer (50 mM Tris-HCl pH 7.5, 250 mM NaCl, 0.1 mM EGTA, 0.1% β-mercaptoethanol). Elute the proteins using a wash buffer containing 20 mM glutathione (pH 7.5). Dialyze into 50 mM Tris-HCl pH 7.5, 0.1 mM EGTA, 150 mM NaCl, 0.5 mM TCEP, 270 mM sucrose, and store at - 70°C. Note: GST-OXSR1 D164A should be stored in 50% glycerol instead of sucrose and kept at -20°C.

#### Baculovirus Expression System (WNK4)

Generation and Infection: Use pFastBac GST-WNK4 (1-469), DU30159, to generate recombinant baculovirus with the Bac-to-Bac system (Invitrogen) as per the manufacturer’s instructions. Infect Spodoptera frugiperda 21 (Sf21) cells grown at 27°C. Harvest the cells 48 h post-infection.

#### Harvesting and Purification

Pellet the cells at 500 x g for 5 min (10 min deceleration). Resuspend in 1/10 of the original culture volume with ice-cold lysis buffer (20 mM HEPES-NaOH pH 7.5, 0.02 mM EGTA, 0.02 mM EDTA, 270 mM sucrose, 0.1% 2-mercaptoethanol, 0.2 mM PMSF, and 1 mM benzamidine). Incubate at 4°C on a roller for 20 min. Purify GST-tagged WNK4 (1-469) as described above and store at -70°C.

### His-SUMO-OXSR1 Expression

#### Expression and Induction

Transform pET His-SUMO-OXSR1 (1-529) D164A, DU55130, into BL21 Codon Plus cells. Induce expression with 150 µM IPTG.

#### Harvesting and Lysis

Resuspend the cell pellet in 20 mL of resuspension buffer (50 mM Tris-HCl pH 7.5, 150 mM NaCl, 2 mM MgCl₂, 20 mM imidazole pH 7.5, 1 mM DTT, Pefabloc (1 mM), and Leupeptin (20 µg/mL)). Lyse cells by adding NaCl (final concentration 300 mM), Triton X-100 (0.2%), and glycerol (5%).

#### Purification

Add 2 mL of equilibrated Nickel agarose (Abcam) resin to the clarified lysate. Incubate at 4°C on a roller for 1 h. Wash with a resuspension buffer containing 5% glycerol and 400 mM NaCl, followed by two washes without imidazole. Elute the His-SUMO-tagged proteins in a wash buffer containing 400 mM imidazole. Dialyze into 50 mM Tris-HCl pH 7.5, 0.1 mM EGTA, 150 mM NaCl, 0.5 mM TCEP, 270 mM sucrose, and store at -70°C.

### Kinase Activity Assays

#### Reaction Setup and Reaction Mixture

Prepare a 50 µL reaction mixture in Tris-based buffer (pH 7.6) containing 50 mM Tris-HCl, 2 mM MgCl₂, and 200 µM ATP. Incubate at 31°C for 20 min to 1 h with agitation at 1000 rpm using a Thermomixer (Thermo Fisher Scientific).

#### Enzymes and Substrates

Use GST-tagged kinase domains of WNK1, WNK2, WNK3, and WNK4, as well as GST or His-SUMO-tagged OXSR1, NRBP1 WT, and NRBP1 Thr232A. The kinase-to-substrate ratio should be 1:10.

#### Stopping the Reaction

Terminate the reaction by adding 12.5 µL of 4X LDS loading buffer with 4 mM β-mercaptoethanol. Heat the samples at 70°C for 5 min to denature the proteins.

#### Phosphorylation Detection

For kinase assays to identify NRBP1 as a substrate of WNK1, perform the assay in the presence of 0.1 mM [γ-32P]ATP (approximately 500 cpm pmol⁻¹) for 2 h.

### Gel Electrophoresis and Detection

#### SDS-PAGE

Resolve the samples on a 10% Bis-Tris gel. Run the gel for 2 h.

#### Gel Staining

Stain the gel using Coomassie InstantBlue for 1 h. Destain with water and scan using an Epson scanner.

#### Gel Drying and Autoradiography

Dry the gel completely using a gel dryer (Bio-Rad). Expose the dried gel to Amersham Hyperfilm for 16 h at -80°C for autoradiography to visualize radiolabeled proteins.

### Identification of NRBP1 Thr232 Phosphosite by LC-MS/MS

#### Kinase Assay, Sample Preparation and Kinase Reaction

Perform kinase assays using kinase-active GST-WNK1 (1-661) and kinase-inactive GST-WNK1 (1-661) V403F. Incubate the reaction for 1h. Process 20% of the reaction mixture for mass spectrometry analysis.

#### Reduction and Alkylation

In a low-binding tube, reduce the proteins by adding 100 mM TCEP (final concentration 5 mM) and incubate at 56°C for 30 min on a thermomixer. Cool the tubes to room temperature. Alkylate non-oxidized sulfhydryl groups with 100 mM IAA (final concentration 20 mM, prepared in 40 mM TEABC, pH 8.5) in the dark at room temperature with shaking at 1000 rpm.

#### Denaturation and Digestion

Denature the proteins with 10 M urea (final concentration 1.5 M, prepared in 40 mM TEABC, pH 8.5). Digest with 200 ng Trypsin Lys-C and incubate at 30°C for 16 h without shaking. Acidify the digested peptides by adding 20% TFA to a final concentration of 1%. Store the acidified peptides at - 20°C until further analysis.

#### Liquid Chromatography and Mass Spectrometry LC Separation

Perform peptide separations using a Thermo Dionex Ultimate 3000 RSLC Nano liquid chromatography system. Use 0.1% formic acid as buffer A and 80% acetonitrile with 0.08% formic acid as buffer B. Load peptides onto a C18 trap column with 3% acetonitrile / 0.1% TFA at a flow rate of 5 μL/min. Separate peptides on an EASY-Spray column (C18, 2 μM, 75 μm x 50 cm) with an integrated nano electrospray emitter at a flow rate of 300 nL/min. Apply a segmented gradient: start at 3% buffer B, increase to 35% over 40 min, then to 95% over 2 min, and hold at 95% for 5 min.

#### MS Analysis

Analyze the eluted peptides using an Orbitrap Lumos (ThermoFisher Scientific) mass spectrometer. Set the spray voltage to 2 kV, RF lens level to 40%, and ion transfer tube temperature to 275°C. Operate in data-dependent mode with 3-second cycles. Perform full scans in the range of 375–1500 m/z with a nominal resolution of 120,000 at 200 m/z, AGC target set to standard, and a maximum injection time of 50 ms. Select the most intense ions above an intensity threshold of 5000 for higher-energy collision dissociation (HCD) fragmentation. Set HCD normalized collision energy to 30%. Acquire data-dependent MS2 scans for charge states 2 to 7 with an isolation width of 1.6 m/z and a 30-second dynamic exclusion duration. Record all MS2 scans in centroid mode with an AGC target set to standard and a maximal fill time of 100 ms.

### Data Processing and Data Analysis

Process the .RAW files using Proteome Discoverer v2.4 (ThermoFisher Scientific) with Mascot v2.6.2 (Matrix Science) as the search engine. Set precursor mass tolerance to 10 ppm and fragment mass tolerance to 0.06 Da. Use an in-house database (MRC_Database_1) with trypsin/P as the protease, allowing a maximum of two missed cleavages. Configure variable modifications to include oxidation and dioxidation of methionine, and phosphorylation of serine, threonine, and tyrosine. Set carbamidomethylation of cysteine as a fixed modification. Utilize ptmRS for scoring phosphosite identification, with a mass tolerance of 0.5 Da and consideration of neutral loss peaks. Confirm phosphorylation site localization only if peptides have a Mascot delta score and ptmRS probability score above 80%.

### Lambda Phosphatase Assay

GFP-NRBP1 was immunoprecipitated from 4 mg of lysate following 30 min of sorbitol treatment. The immunoprecipitated protein was resuspended in 100 μL of assay buffer containing 50 mM HEPES (pH 7.5), 100 mM NaCl, 2 mM DTT, 0.01% Brij, and 2 mM MgCl₂, and then divided equally into two tubes. To one tube, approximately 2 μg of Lambda phosphatase (DSTT) was added. Both tubes were incubated at 30°C for 1 h with gentle agitation at 1200 rpm. The reaction was stopped by adding 10 μL of 4X LDS loading dye containing 4% β-mercaptoethanol and heating at 90°C for 5 min.

### Generation of Stable Cell Lines Using CRISPR/Cas9

For GFP knock-ins, HEK-293 cells were transfected with 1 μg of each plasmid vector encoding antisense and sense guide RNAs targeting NRBP1 or WNK1, along with 3 μg of a GFP donor plasmid. For the generation of the Bromotag NRBP1 knock-in cell line, 2 μg of the donor plasmid and 1 μg of the antisense guide RNA were used. Transfection was carried out for 24 h, after which cells were selected with 2 μg/mL puromycin (Sigma, P9620) for 48 h. The cells were then allowed to recover in fresh media until they reached confluency. Fluorescence-activated cell sorting (FACS) was employed to isolate GFP-positive cells into single wells of two 96-well plates containing conditioned media. Surviving cell clones were expanded, and the successful knock-in of GFP or Bromotag to the N-terminus was verified using western blotting, immunoprecipitation, and DNA sequencing. The NRBP1 KO discussed in SFig 7 were obtained while selecting the Bromotag NRBP1 KI clones. To knock out NRBP1 from the GFP-NRBP1 knock-in HEK-293 cell line, a CRISPR construct targeting GFP was used. GFP-negative cells were then selected for further analysis.

### Generation of Stable Cell Lines by Retroviral Transduction

To generate stable cell lines, the following retroviral components were prepared: 6 μg of pBABE vectors (pBABE puro HA NRBP1 [DU77418] or pBABE puro HA NRBP1 Thr232A [DU77419]), 3.2 μg of pCMV5-GAG/POL, 2.8 μg of pCMV5-VSV-G, and 20 μl of PEI (1 mg/ml). These components were added to 1 mL of Opti-MEM (31985–062, GIBCO). The mixture was vortexed gently and incubated at room temperature for 20 min. The transfection mixture was then added dropwise to HEK-293-FT cells (at approximately 70% confluency) growing in a 10 cm diameter cell culture dish. After 24 h, the media was changed to a fresh medium. Following an additional 24 h, the media containing retroviruses was collected and filtered through a 0.22 μm sterile filter. Target cells (at 60% confluency) were then transduced with the viral supernatant in the presence of 8 μg/mL hexadimethrine bromide (also known as polybrene, H9268, Sigma) for 24 h. After transduction, the media was replaced with fresh media containing 2.5 μg/mL puromycin (P9620, Sigma) to select for successfully transduced cells. Selection with puromycin was carried out for 48 h, after which the cells were maintained in fresh culture media.

### Activation of the WNK1 Signaling Pathway by Hypertonic Stress

To activate the WNK1 signaling pathway, cells were exposed to complete DMEM containing 0.5 M sorbitol (diluted in 1X PBS) for 5-30 min, as previously described (Zagórska et al., 2007). Prior to this, cells were pretreated for 20 min with either the pan-WNK inhibitor WNK463 (DSTT) or the WNK1/3-specific inhibitor WNK C11/12 (DSTT) at a concentration of 10 μM. Following pretreatment, cells were incubated in the respective treatment media containing the inhibitors.

### Biotin Proximity Labeling with TurboID

To activate the biotin-ligase activity of TurboID, exogenous biotin was added to the culture medium as previously described (*45, 79*). Briefly, the growth media was aspirated from cell culture dishes and replaced with fresh DMEM supplemented with 10% (by vol) FBS, 2 mM L-glutamine, 500 μM biotin (Sigma Aldrich, B4501), and optionally, 0.5 M sorbitol (Sigma Aldrich, S1876). After 5 min, the media was aspirated, and cells were washed rapidly with ice-cold 1X PBS three times. Cells were then lysed in buffer containing 50 mM Tris, pH 7.5, 0.27 M sucrose,150 mM NaCl, 1 mM EDTA, 1 mM EGTA, 1% (by vol) NP-40, 1 mM sodium vanadate (NaVO4), 5 mM sodium pyrophosphate (Na_4_P_2_O_7_), 50 mM sodium fluoride (NaF); 10 mM β-glycerophosphate; and a 1× EDTA-free protease inhibitor cocktail [Merck, 11873580001]). Lysates were incubated on ice for 10 min and then clarified by centrifugation at 30,000 g for 10 min at 4 °C. The supernatant (soluble cell extract) was collected in a fresh Eppendorf tube and snap-frozen in liquid nitrogen and stored at −80 °C. The protein concentration was estimated by the Bradford protein assay (Pierce, 23236). We employed a TurboID variant known for its substantially improved biotinylation kinetics, reducing labeling times compared to BioID, and eliminating the need for hydrogen peroxide treatment to induce activation (*45, 79*).

### Streptavidin Enrichment of Biotinylated Proteins from Cell Lysate for LC-MS Analysis

For LC-MS analysis, 3.5 mg of cell lysate was used for immunoprecipitation with 25 µl of streptavidin magnetic beads (Pierce 88817) for 2 h at 4°C with gentle rotation. The beads were washed four times with cold 1X PBS containing 0.05% (by vol) NP-40 by inverting the tubes 20 times. The beads were resuspended in 0.5 ml of 50 mm Tris pH 7.5, immobilized on a magnetic rack, and 100 µL of the bead slurry was used for western blotting.

For reduction, the bead pellet was resuspended in 50 µL of 5% SDS-TEABC (12.5 µL of 20% SDS and 40 µL of 50 mM TEABC) and 5 mM TCEP (2.5 µl of 100 mM stock prepared in 50 mM TEAB, pH 8.5), and incubated at 60°C for 30 min at 1200 rpm on a thermomixer with lid. After cooling to room temperature, 20 mM iodoacetamide (IAA) (10 µL of 100 mM stock made in MQ water) was added and incubated at room temperature for 30 min at 1200 rpm on a thermomixer. Alkylation was quenched with 2.5 mM TCEP (1.25 µL per tube) for 10 min at room temperature, and the samples were stored at -80°C. Next day, the samples were thawed and 6µl of 27.5% phosphoric acid was added and briefly vortexed. The beads were immobilized, the supernatant was collected in a fresh low binding tube and the volume was made to 0.5 ml with the binding buffer (5 ml of 1 M TEAB + 45 ml methanol. Trypsinization and peptide elution were performed using the S-Trap micro column as per the manufacturer’s protocol (Protifi, C02-micro-80). Briefly, 20 µg of Trypsin/LysC mix (MS grade, Pierce 1863520) was reconstituted in 200 µL of 100 mM TEAB to a stock solution of 0.1 µg/mL. 20 µL of this solution (2 µg) was loaded onto the trap column and incubated at 47°C for 2 h on a thermomixer with lid and no shaking. Digested peptides were sequentially eluted using 40 µL of Elution Buffer 1 (50 mM TEAB in water, pH 8.5), Elution Buffer 2 (0.2% formic acid in water), and Elution Buffer 3 (50% acetonitrile in water) by centrifugation at 1500 x g for 1 minute. The eluted peptides were pooled, dried in a SpeedVac for 2 h, and resuspended in 20 µL of 5% formic acid in water.

Peptides were analyzed by injecting 200 ng onto a Vanquish Neo UHPLC System operating in trap and elute mode coupled with an Orbitrap Astral Mass Spectrometer (Thermo Fisher Scientific). Peptides were loaded onto a PepMap Neo Trap Cartridge (Thermo Fisher Scientific #174500) and analyzed on a C18 EASY-Spray HPLC Column (Thermo Fisher Scientific #ES906) with a 11.8 minute gradient from 1% to 55% Buffer B (Buffer A: 0.1% formic acid in water; Buffer B: 0.08% formic acid in 80:20 acetonitrile:water, 0.7 min at 1.8 µL/min from 1% to 4% B, 0.3 min at 1.8 µL/min from 4% to 8% B, 6.7 min at 1.8 µL/min from 8% to 22.5% B, 3.7 min at 1.8 µL/min from 22.5% to 35% B, 0.4 min at 2.5 µL/min from 35% to 55% B). Eluted peptides were analyzed using data-independent acquisition mode on the mass spectrometer.

### Streptavidin Enrichment of Biotinylated Proteins for Western Blot Analysis

For Western blot analysis, 2 mg of lysate was incubated with 20 µL of streptavidin magnetic beads for 2 h at 4°C with gentle rotation. The beads were washed thrice with cold 1X PBS containing 0.05% (by vol) NP-40 and then eluted by boiling the beads for 10 min in 50 µL of 2X NuPAGE LDS buffer containing 2 mM biotin.

### Immunoprecipitation of GFP NRBP1 for Mass Spectrometry

To identify the direct interaction with WNK’s and other sorbitol stimulated interactors, GFP-NRBP1 was immunoprecipitated from the GFP-NRBP1 KI HEK293 cells with WT HEK293 serving as negative and bead control. The experiment involved 5 technical replicates for each of the 4 groups namely, GFP-NRBP1 KI HEK293 and WT HEK293 with and without 0.5 M sorbitol for 30 min. Around 4 mg of the lysate in 1 ml lysis buffer (20 mM HEPES pH7.5,150 mM NaCl, 1 mM EDTA, 1 mM EGTA, 0.5% (by vol) NP-40, 1 mM NaVO4, 5 mM Na pyrophosphate, 50 mM Sodium fluoride, 10 mM β-Glycerophosphate and a protease inhibitor mini tab) was immunoprecipitated using GFP magnetic beads (GFP-Trap® Magnetic Particles M-270 Kit). 25 μl of magnetic bead slurry per IP was used and the immunoprecipitation was carried out at 4°C for 2 h. The beads were then immobilized on a magnetic rack and 50 μl of the supernatant was collected to check for the depletion of GFP-NRBP1 The beads were washed by inverting the tubes 20 times with Wash Buffer 1 (lysis buffer with 0.1% (by vol) NP-40) twice; Wash Buffer 2 (lysis buffer without NP-40) twice; and Wash Buffer 3 (20 mM HEPES, pH 7.5) to remove salts. The beads were resuspended in 0.5 ml of 20 mM HEPES, and 10% (50 µL) of the bead slurry was reserved for western blotting. The reduction, alkylation, trypsinization and elution steps were same as that for streptavidin enrichment of biotinylated proteins.

Peptides were analyzed by injecting 2 µg onto an UltiMate 3000 RSLCnano System coupled with an Orbitrap Fusion Lumos Tribrid Mass Spectrometer (Thermo Fisher Scientific). Peptides were loaded onto an Acclaim PepMap trap column (Thermo Fisher Scientific #164750) and analyzed on a C18 EASY-Spray HPLC Column (Thermo Fisher Scientific #ES903) with a 120-minute linear gradient from 3% to 35% Buffer B (Buffer A: 0.1% formic acid in water; Buffer B: 0.08% formic acid in 80:20 acetonitrile:water). Eluted peptides were analyzed using data-independent acquisition mode on the mass spectrometer.

### Data Analysis

Peptides were searched against the UniProt SwissProt Human database (released on 05/10/2021) using DiaNN (v1.8.1) in library-free mode. Statistical analysis was performed with Python (v3.9.0) and the following packages: pandas (v1.3.3), numpy (v1.19.0), sklearn (v1.0), scipy (v1.7.1), rpy2 (v3.4.5), Plotnine (v0.7.1), and Plotly (v5.8.2). Additionally, R (v4.1.3) and the Limma package (v3.50.1) were used for analysis. Protein groups with fewer than 2 proteotypic peptides or quantified in fewer than 3 replicates were excluded from further analysis. Missing values were imputed using a Gaussian distribution centered on the median, with a downshift of 1.8 and a width of 0.3 relative to the standard deviation, and protein intensities were median-normalized. Protein regulation was assessed using LIMMA, with p-values adjusted by the Benjamini-Hochberg multiple hypothesis correction. Proteins were considered significantly regulated if their corrected p-value was less than 0.05 and their fold change was either greater than 1.5 or less than 1/1.5.

### AlphaFold 3 Modelling

Modelling of structures of individual proteins and protein complexes was performed using AlphaFold 3 (*60*) with primary sequences obtained from UniProt: NRBP1 (Q9UHY1), SPAK (Q9UEW8), TSC22D2 (O75157), TSC22D4 (Q9Y3Q8), WNK1 (Q9H4A3), WNK2 (Q9Y3S1), WNK3 (Q9BYP7), WNK4 (Q96J92). The full-length sequences were used to predict the structures of monomers of NRBP1, WNK1, WNK2, WNK3, WNK4, TSC22D2, TSC22D4, homo- and heterodimers of TSC22D2 and TSC22D4, homodimer of NRBP1, complexes of NRBP1 with TSC22D2 or TSC22D4, as well as WNK1 in complex with NRBP1, WNK1 in complex with SPAK and NRBP1, WNK1 in complex with SPAK, NRBP1 and TSC22D4 homodimer. Detailed intermolecular interactions for models of NRBP1 in complex with TSC22D2 or TSC22D4 were analyzed with BIOVIA Discovery Studio Visualizer 2024 using the Non-bond Interaction Monitor. All structures were visualized using PyMOL 3.0

### List of constructs generated

The constructs are available on requests made to Medical Research Council (MRC) – Phosphorylation and Ubiquitylation Unit (PPU) Reagents webpage https://www.ppu.mrc.ac.uk/ and the unique identifier (DU) numbers indicated below provide direct links to the cloning strategy and sequence information.

| DU Number | Construct Name |
| --- | --- |
| **For Transient Transfection studies** |  |
| DU 29702 | pBABED puro FLAG-TurboID-aGFP<sub>6M</sub> |
| DU68362 | pcDNA5D FRT/TO HA NRBP1 |
| DU72784 | pcDNA5D FRT/TO HA NRBP1 L476A |
| DU72766 | pcDNA5D FRT/TO HA NRBP1 E484A |
| DU72783 | pcDNA5D FRT/TO HA NRBP1 E492A |
| DU72782 | pcDNA5D FRT/TO HA NRBP1 E484A + E492A |
| DU68365 | pcDNA5D FRT/TO HANRBP1 Thr232A |
| DU72496 | pcDNA5D FRT/TO GFP TSC22D2 |
| DU72456 | pcDNA5D FRT/TO GFP TSC22D4 |
| DU80387 | pCMV5D Flag TSC22D2 |
| DU76241 | pCMV5D FLAG TSC22D2 R161A |
| DU80408 | pCMV5D FLAG TSC22D2 R161A F162A |
| DU80409 | pCMV5D FLAG TSC22D2 R161E F162A |
| DU80407 | pCMV5D FLAG TSC22D2 R179A |
| DU80407 | pCMV5D FLAG TSC22D2 R179E |
| DU80449 | pCMV5D FLAG TSC22D2 R161E F162AR179E |
| DU61990 | pCMV5D FLAG TSC22D2 M1-I690 |
| DU68295 | pCMV5D HA TSC22D2 |
| DU77694 | pCMV5D Flag TSC22D4 |
| DU61975 | pCMV5D FLAG TSC22D4 M1-N320 |
| **For protein expression** |  |
| DU43603 | GST-WNK1 |
| DU49486 | GST-WNK2 |
| DU 4409 | GST-WNK3 |
| DU2121 | GST-WNK4 |
| DU2961 | GST-OXSR1 D164A |
| DU55144 | pet15 6HIS SUMO OXSR1 WT FL 1-527 |
| DU68346 | pGEX6P1 NRBP1 |
| DU68840 | pGEX6P1 NRBP1 Thr232A |
| DU72490 | pET15b 6HIS SUMO NRBP1 |
| **For CRISPR mediated knockout and knock-in studies** |  |
| DU69224 | pMK NRBP1 Nter KI donor- donor puro Bromotag |
| DU64963 | pX459 NRBP1 Nter KI sgG1 (guide + Cas9+ puro) (used for both GFP NRBP1 KI and Bromotag NRBP1 KI) |
| DU69003 | pMK-RQ NRBP1 Nter GFP donor |
| DU29702 | pBABED puro FLAG-TurboID-aGFP<sub>6M</sub> |
| DU57149 | pWNK1 Nter GFP IRES2 APEX2 V5 donor |
| DU52054 | pBabeD P U6 WNK1 Nter KI Sense A |
| DU52062 | pBabeD P U6 WNK1 Nter KI Antisense A |
| **For retroviral transfections** |  |
| DU77418 | pBabeD puro HA NRBP1 |
| DU77419 | pBabeD Puro HA NRBP1 Thr232A |

## Acknowledgments

We thank Gerrit Daubner for generation of the GFP-WNK1 HEK293 cell line and the technical support of the MRC Protein Phosphorylation and Ubiquitylation Unit (PPU) including the MRC-PPU DNA sequencing service (coordinated by G. Hunter), the MRC-PPU tissue culture team (coordinated by E. Allen), the MRC-PPU MS facility team, and the MRC-PPU Reagents and Services team (coordinated by J. Hastie).

---

**Supplementary Fig 1:**
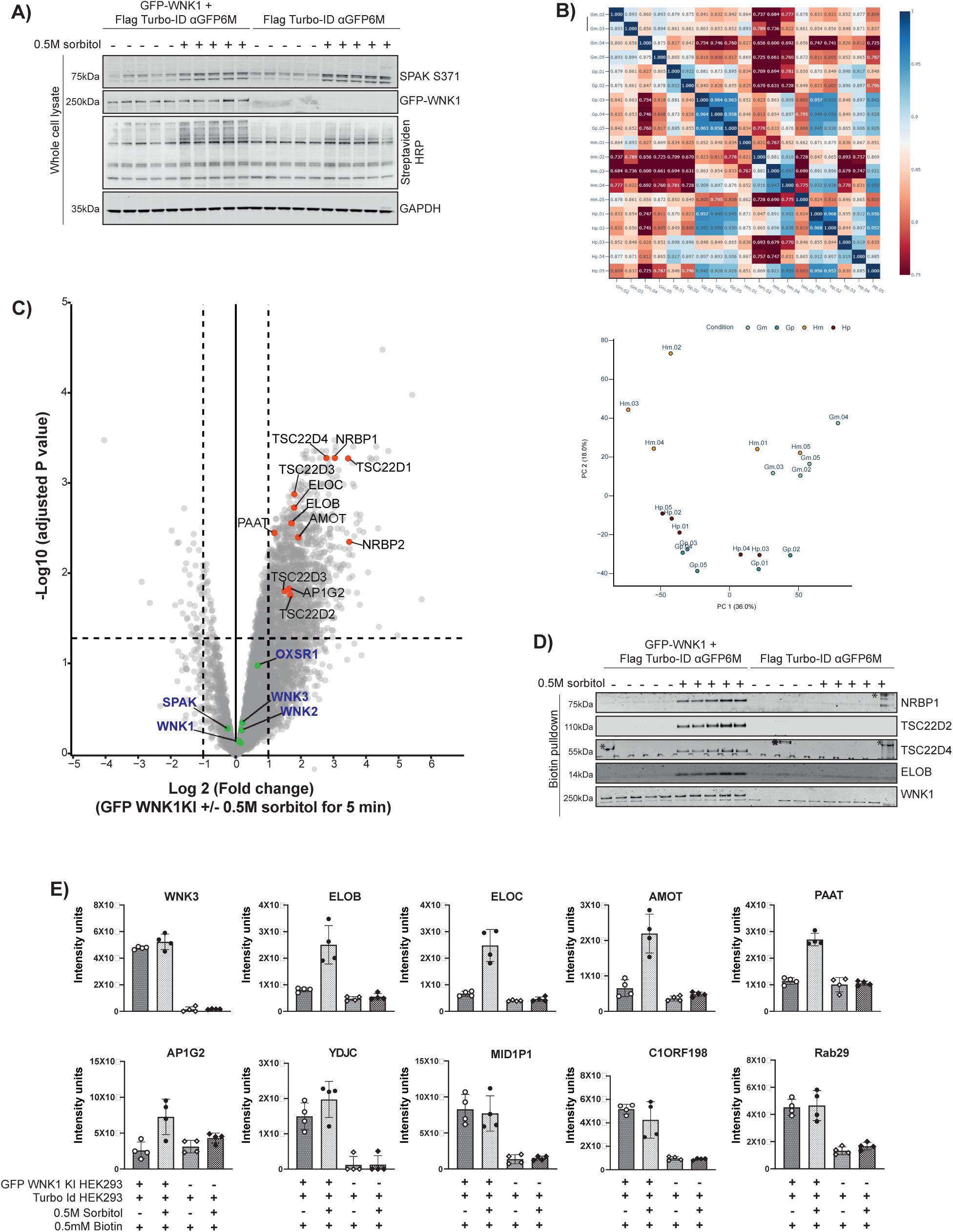
Analysis of WNK Pathway Activation and Protein Biotinylation Upon Hypertonic Stress. **(A)** Representative immunoblots showing the activation of the WNK signaling pathway (pSPAK blot) and enhanced protein biotinylation (streptavidin-HRP blot) in response to 0.5 M sorbitol treatment for 5 minutes in the presence of 0.5 mM biotin. Data represent biological replicates (n = 5 for each treatment group). **(B) Top Panel:** Heatmap displaying the Pearson correlation coefficients between imputed, median-normalized protein intensities across samples, illustrating overall consistency in replicate samples. **Bottom Panel:** Principal component analysis (PCA) plot showing the clustering of different treatment groups. Variability within groups is noted, potentially due to the short treatment time, which may limit the magnitude of large-scale proteomic changes. Note: Gm-GFP WNK1 cells without sorbitol, Gp-GFP WNK1 cells with sorbitol, Hm-WT WNK1 cells without sorbitol and Hp-WT WNK1 cells with sorbitol.**(C)** Volcano plots highlighting proteins enriched in GFP-WNK1 KI HEK293 cells expressing FLAG-TurboID-aGFP6M treated with 0.5 mM biotin, with or without 0.5 M sorbitol. Protein intensities were normalized, imputed using a Gaussian distribution, and subjected to Benjamini-Hochberg correction for multiple hypothesis testing. Proteins with ≥2-fold enrichment and adjusted P-values <0.05 are annotated on the volcano plots, visualized using the Curtain tool. Proteins highlighted in green include the TurboID bait (WNK1) and its known interactors (WNK2, WNK3, SPAK, and OXSR1). Proteins highlighted in red are sorbitol-specific differential interactors of WNK1. **(D)** Western blot analysis of streptavidin pull-down fractions from GFP-WNK1 TurboID cells. Biotinylation of NRBP1, TSC22D2, TSC22D4, and ELOB is observed exclusively following hypertonic stress, validating their stress-specific enrichment. **(E)** Box plots depicting the median protein intensities of significantly enriched hits from the volcano plot in (C) and from Figure 1(D).

**Supplementary Figure 2.**
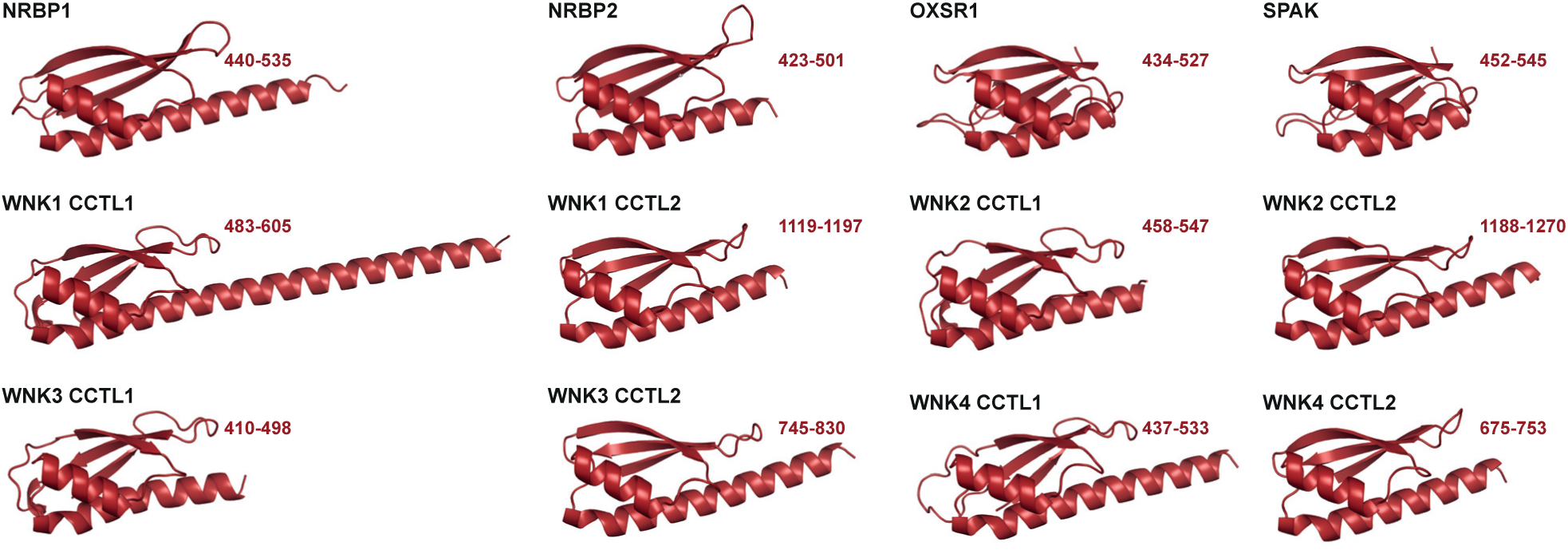
Structural Analysis of CCT and CCTL Domains in Signaling Proteins. Structures of the CCT and CCTL domains of NRBP1, NRBP2, OXSR1, SPAK, WNK1, WNK2, WNK3, and WNK4, as derived from full-length AlphaFold 3 models.

**Supplementary Figure 3.**
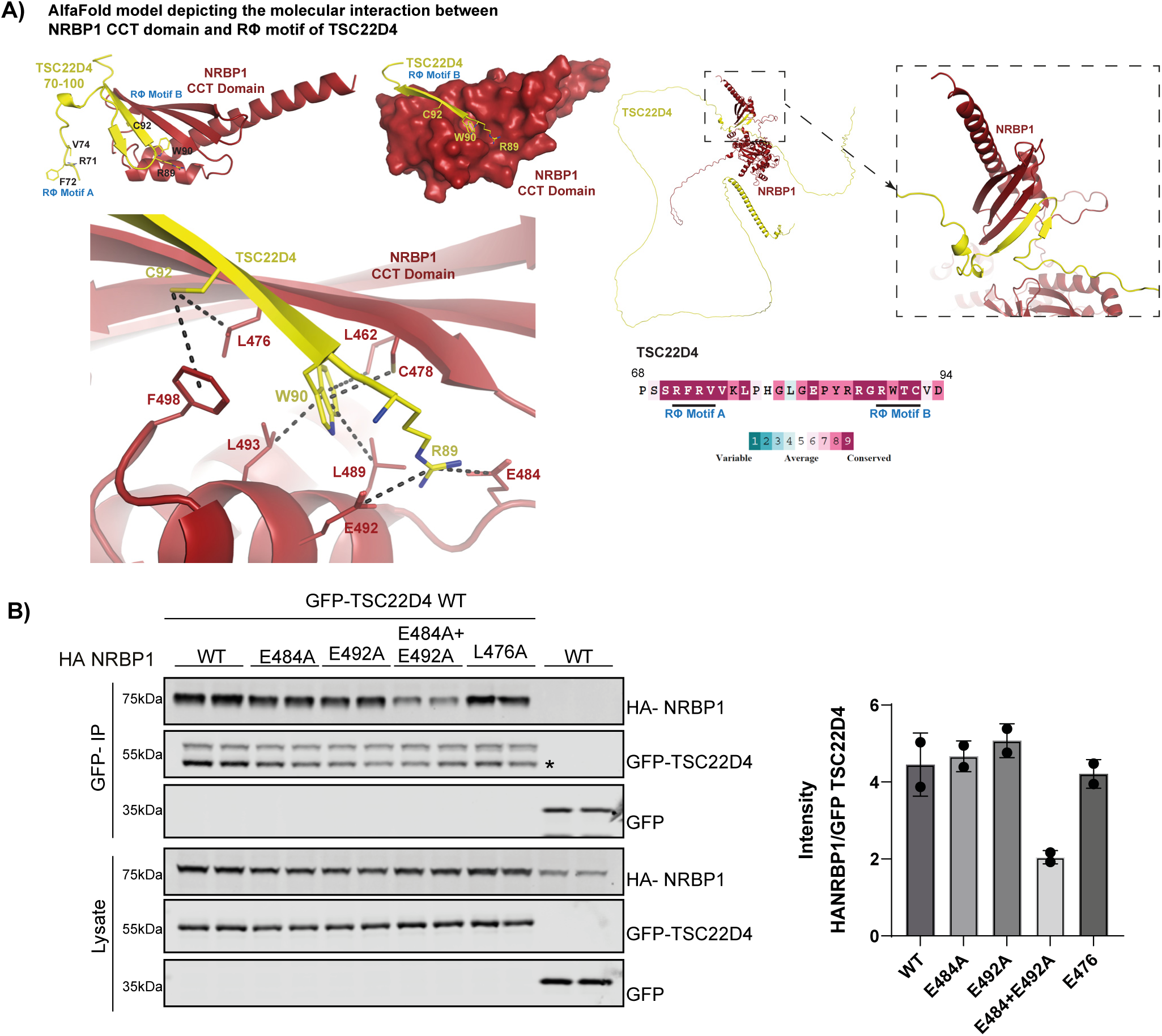
Interaction Analysis Between the CCT Domain of NRBP1 and the RΦ Motif B of TSC22D4. **(A)** Detailed model of the interaction between the CCT domain of NRBP1 and the RΦ motif B of TSC22D4, extracted from a full-length AlphaFold 3 prediction. Trp90 of TSC22D4’s RΦ motif B binds within a hydrophobic pocket of the CCT domain of NRBP1, formed by Leu462, Leu464, Leu476, Cys478, Leu489, Leu493, Leu496, and Phe498. Additional hydrophobic interactions are observed between Cys92 of TSC22D4 and Leu476 and Phe498 of NRBP1. Salt bridges are formed between Arg89 of TSC22D4 and Glu484 and Glu492 of NRBP1. ConSurf analysis (*80*) shows high conservation of RΦ motifs A and B in TSC22D4. **(B)** Co-immunoprecipitation of GFP-TSC22D4 WT and HA-NRBP1 mutants (E484A, E492A, and E476A) in HEK 293 cells after 36 h of co-transfection. Interaction was studied by GFP immunoprecipitation, followed by western blotting. **Right**: Densitometric analysis of the western blots. Data are representative of n=3 experiments with two technical replicates for each condition, analyzed by one-way ANOVA (**P < 0.01, ***P < 0.001, and ****P < 0.0001).

**Supplementary Figure 4.**
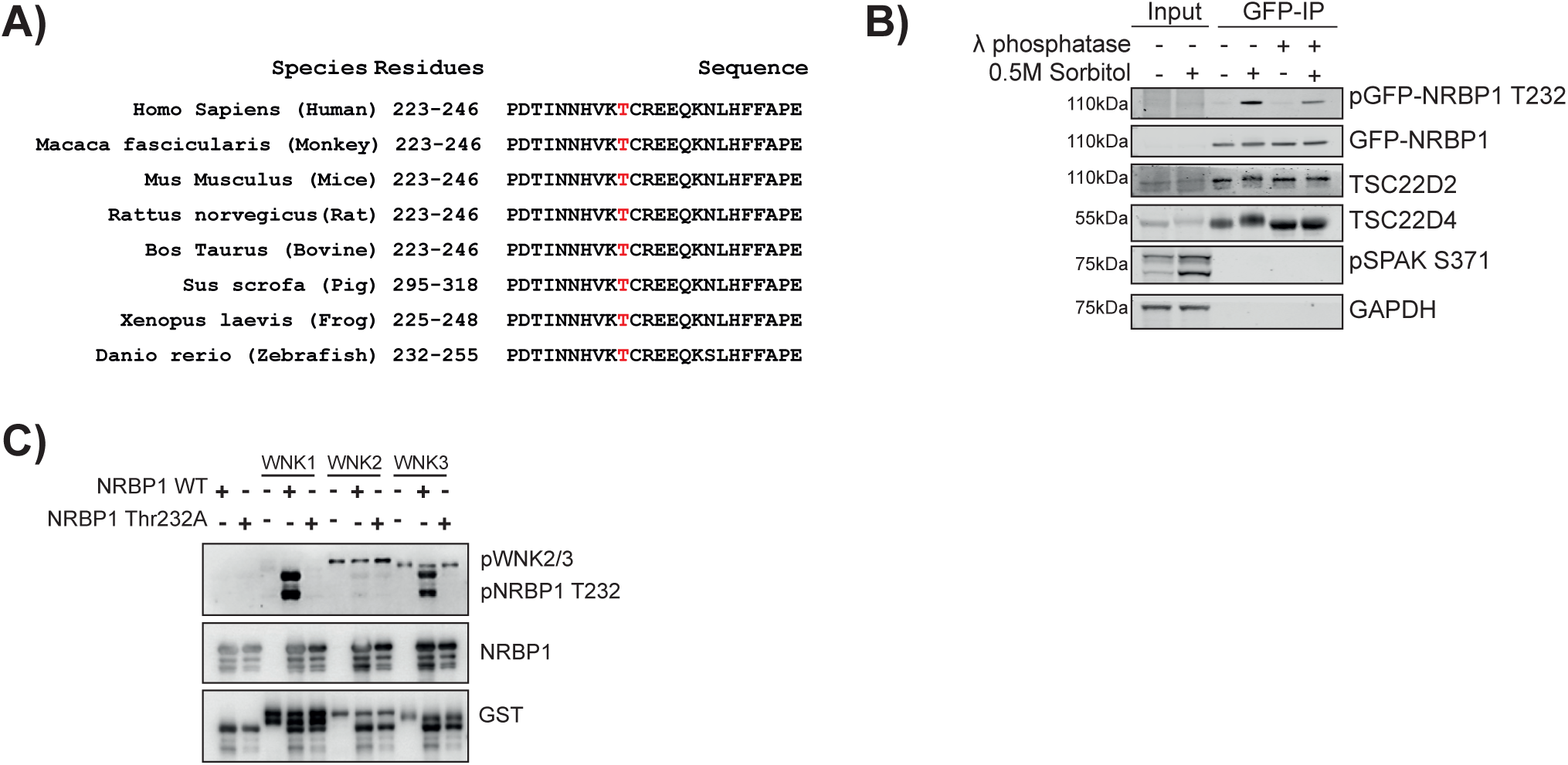
Phosphorylation of NRBP1 and Its Interaction with TSC22D2/4. **(A)** Sequence alignment of the activation T-loop of NRBP1 from different species, performed using the MUSCLE tool (*81*), showing the evolutionary conservation of Thr232 (highlighted in red, numbering based on human sequence). **(B)** GFP-NRBP1-TSC22D2/4 complex was immunoprecipitated post-sorbitol treatment, and the immunoprecipitate was subjected to λ-phosphatase treatment to validate whether the observed TSC22D4 upshift is due to phosphorylation. **(C)** *In vitro* kinase assay with GST-tagged WNK1, WNK2, WNK3 kinases, and GST-tagged NRBP1 (WT & Thr232A mutant), showing that NRBP1 Thr232 is phosphorylated by WNK1 and WNK3. Results from n=2 are shown as a representative experiment.

**Supplementary Figure 5.**
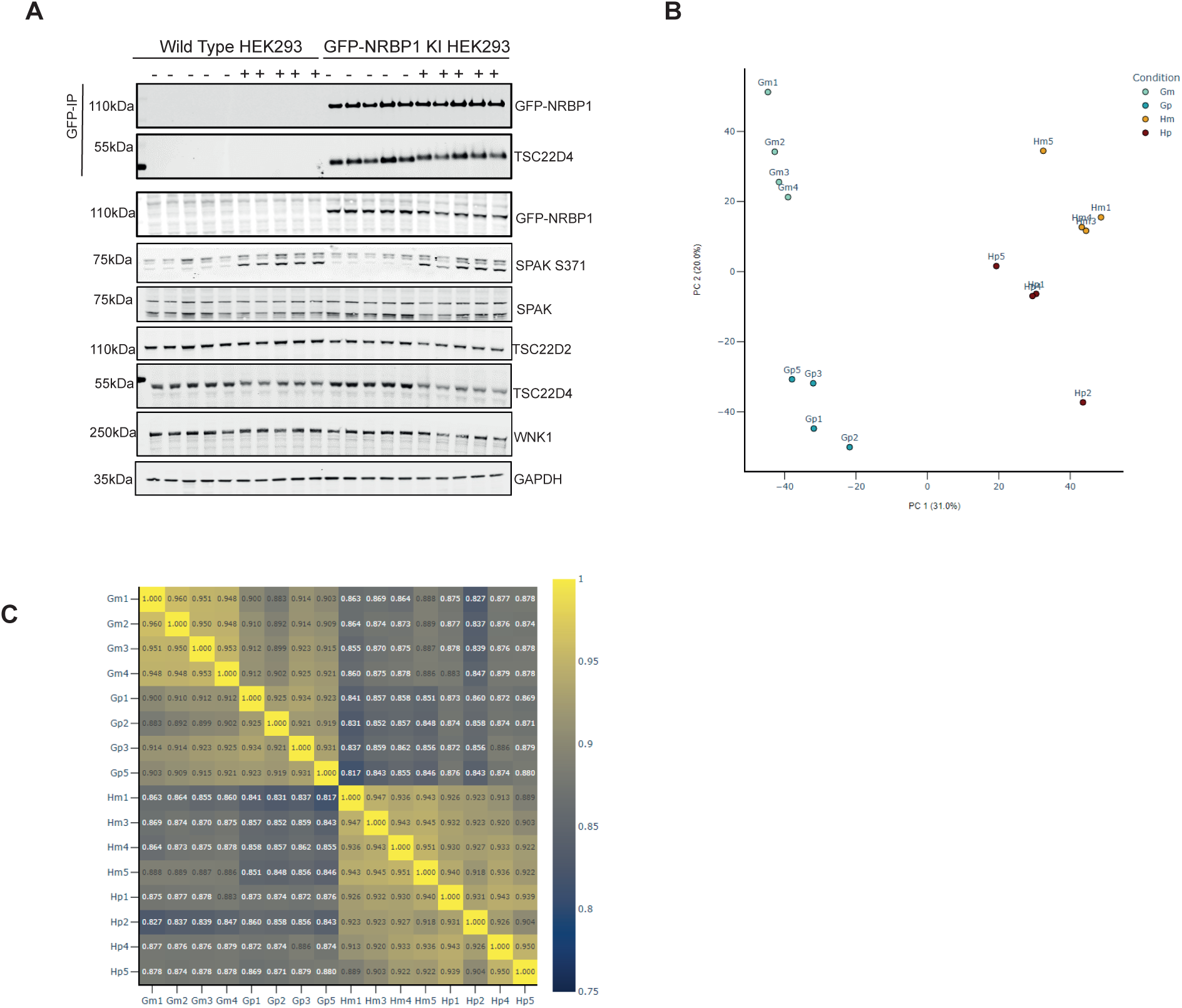
Validation and Analysis of NRBP1 Interactors. (A) Immunoblots showing the activation of the WNK pathway and successful immunoprecipitation of GFP-NRBP1 for the mass spectrometry (MS) experiment described in Figure 5. N=5 technical replicates for each group. (B) Principal Component Analysis (PCA) plot showing the clustering of different treatment groups. (C) Heatmap displaying the Pearson correlation coefficients between the imputed, median-normalized protein intensities across each sample. Note: Gm-GFP NRBP1 cells without sorbitol, Gp-GFP NRBP1 cells with sorbitol, Hm-WT NRBP1 cells without sorbitol and Hp-WT NRBP1 cells with sorbitol.

**Supplementary Figure 6.**
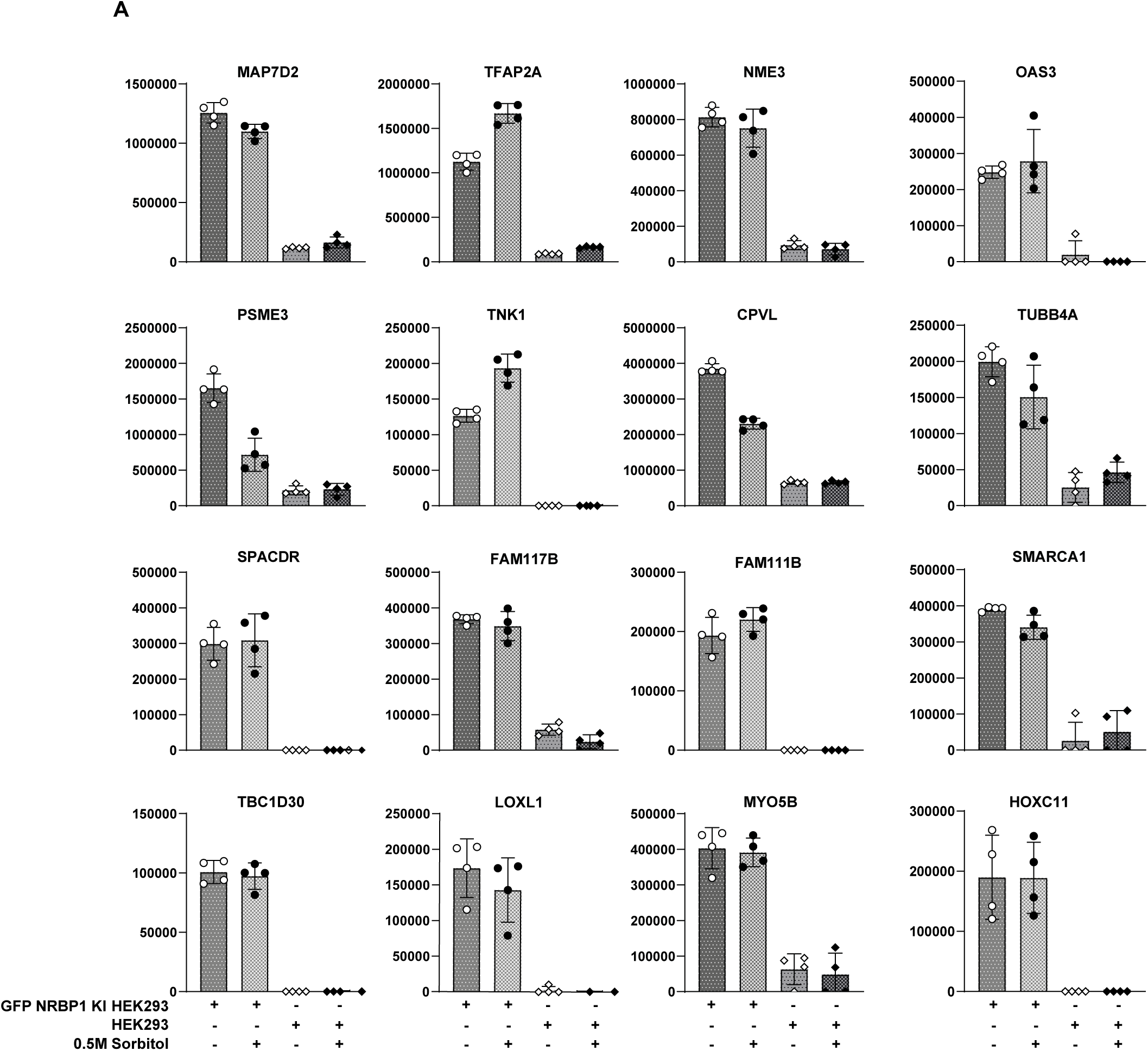
Analysis of NRBP1 Interactors. Box plots showing the protein median intensities of selected NRBP1 interactors identified from the volcano plot in Figure 5B.

**Supplementary Figure 7.**
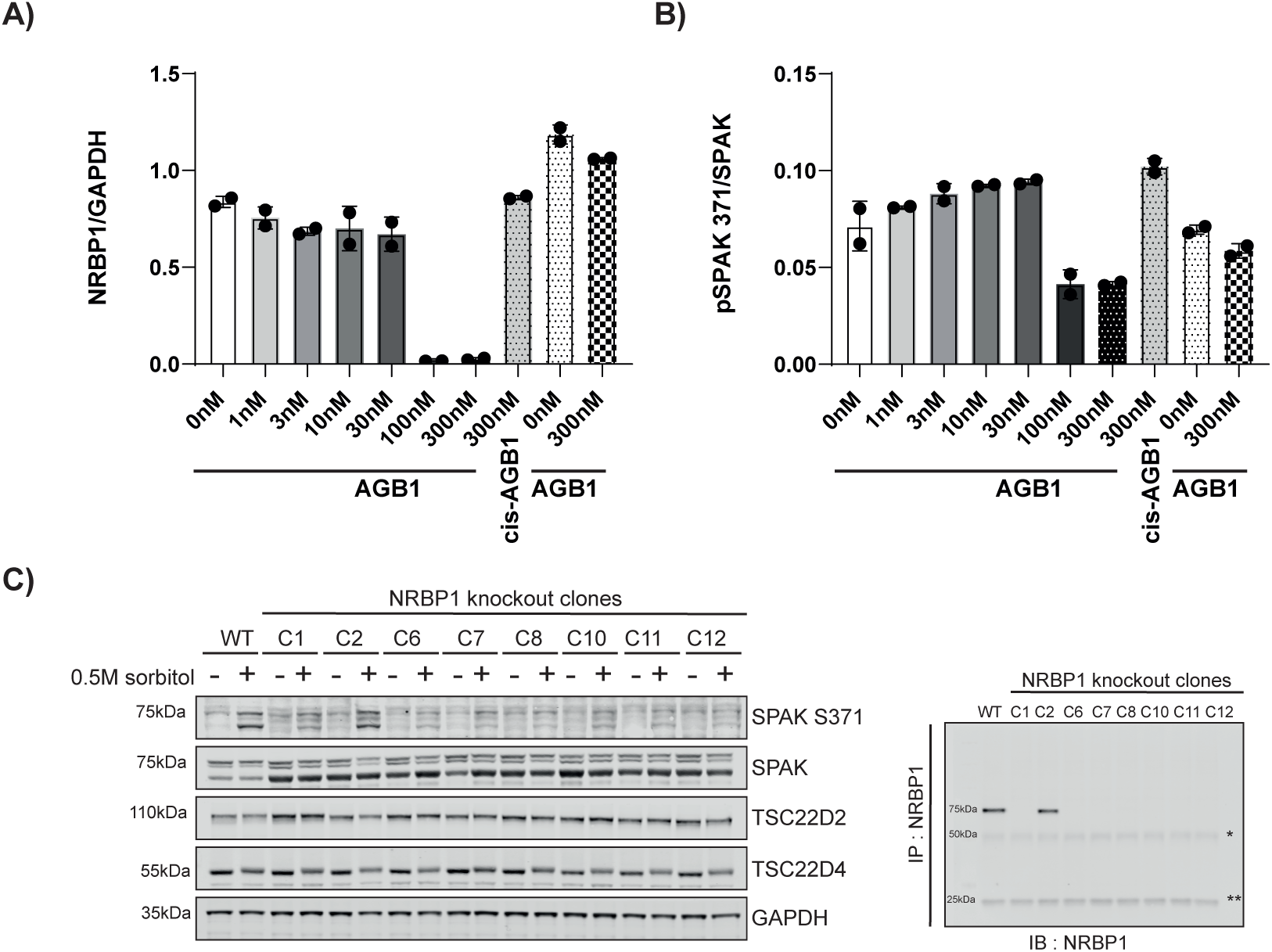
Analysis of NRBP1 and SPAK Phosphorylation. **(A & B)** Densitometric analysis of NRBP1 and pSPAK levels, respectively, from the immunoblot in Figure 7A. **(C) Left Panel**: Effect of NRBP1 knockout (KO) on SPAK phosphorylation was analyzed by immunoblotting following 0.5 M sorbitol treatment for 30 min in 7 NRBP1 knockout HEK293 clones. **Right Panel**: Validation of NRBP1 knockout by endogenous immunoprecipitation and immunoblotting.

**Supplementary Figure 8.**
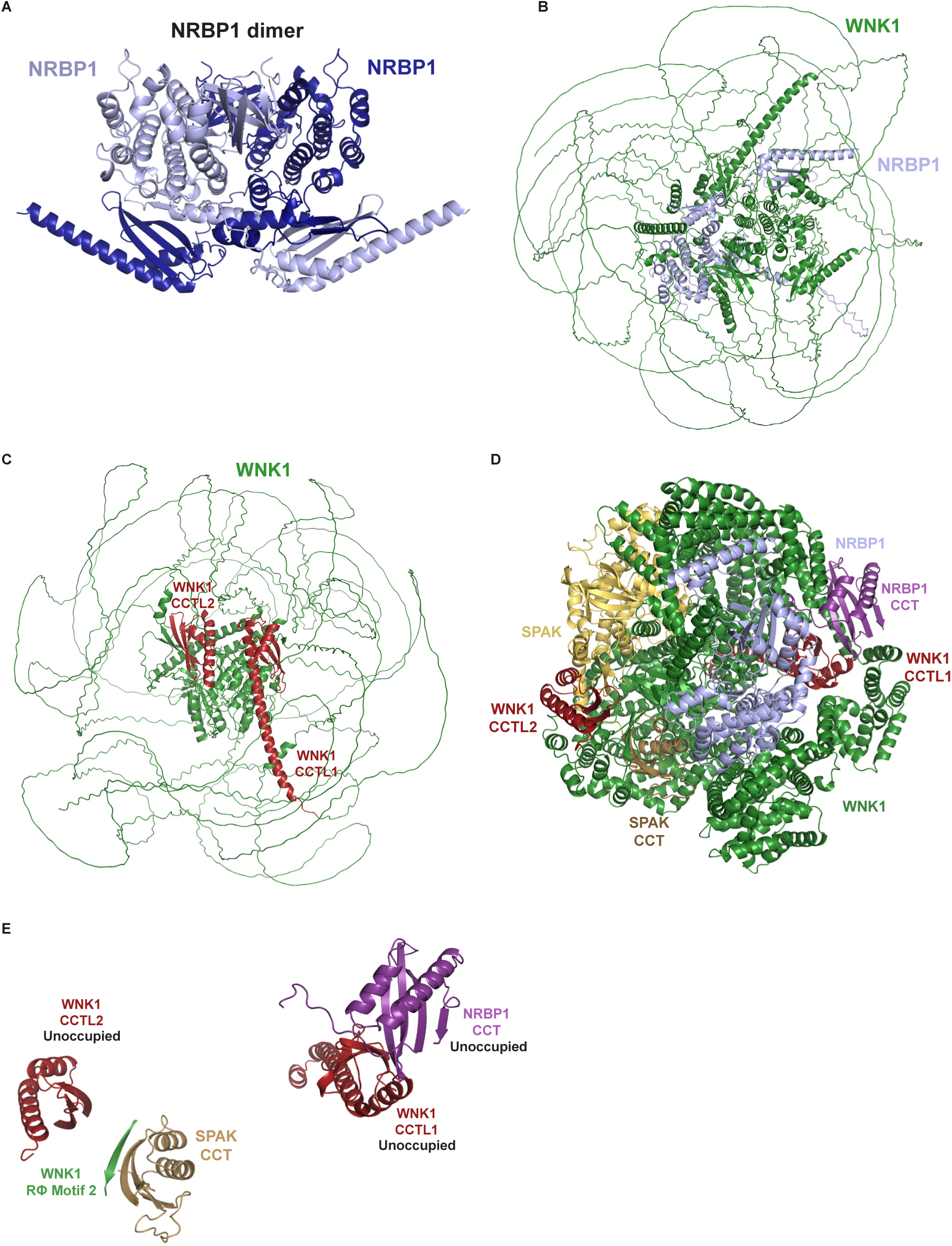
AlphaFold 3 Models of NRBP1 and WNK1 Complexes. **(A)** AlphaFold 3 model of the NRBP1 homodimer. **(B)** AlphaFold 3 model of WNK1 in complex with NRBP1. **(C)** AlphaFold 3 model of full-length WNK1 with CCTL domains highlighted in red. **(D)** AlphaFold 3 model of WNK1 in complex with SPAK and NRBP1. **(E)** As in (D), but only the CCT domains and RΦ motifs are shown.

